# Aged-senescent cells contribute to impaired heart regeneration

**DOI:** 10.1101/397216

**Authors:** Fiona C. Lewis-McDougall, Prashant J. Ruchaya, Eva Domenjo-Vila, Tze Shin Teoh, Larissa Prata, Beverley J. Cottle, James E. Clark, Prakash P. Punjabi, Wael Awad, Daniele Torella, Tamara Tchkonia, James L. Kirkland, Georgina M. Ellison-Hughes

## Abstract

Aging leads to increased cellular senescence and is associated with decreased potency of tissue-specific stem/progenitor cells. Here we have done an extensive analysis of cardiac progenitor cells (CPCs) isolated from human subjects with cardiovascular disease (n=119), aged 32-86 years. In aged subjects (>74 years old) over half of CPCs are senescent (p16^INK4A^, SA-β-gal, DNA damage γH2AX, telomere length, Senescence-Associated Secretory Phenotype (SASP)), unable to replicate, differentiate, regenerate or restore cardiac function following transplantation into the infarcted heart. SASP factors secreted by senescent CPCs renders otherwise healthy CPCs to senescence. Elimination of senescent CPCs using senolytics abrogates the SASP and its debilitative effect *in vitro*. Global elimination of senescent cells in aged mice (INK-ATTAC or wildtype mice treated with D+Q senolytics) *in vivo* activates resident CPCs (0.23±0.06% vs. 0.01±0.01% vehicle; p<0.05) and increased the number of small, proliferating Ki67-, EdU-positive cardiomyocytes (0.25±0.07% vs. 0.03±0.03% vehicle; p<0.05). Therapeutic approaches that eliminate senescent cells may alleviate cardiac deterioration with aging and rejuvenate the regenerative capacity of the heart.

## Introduction

Ageing is the greatest risk factor for many life-threatening disorders, including cardiovascular diseases, neurodegenerative diseases, cancer, and metabolic syndromes [1]. Although long-term exposure to known cardiovascular risk factors strongly drives the development of cardiovascular pathologies, intrinsic cardiac aging is considered to highly influence the pathogenesis of heart disease [2]. However, the fields of the biology of aging and cardiovascular disease have been studied separately, and only recently their intersection has begun to receive the appropriate attention.

Aging leads to increased cellular senescence in a number of tissues and work suggests senescent cell burden can be dramatically increased in various tissues and organs with chronological ageing or in models of progeria [3-5]. Cellular senescence is associated with increased expression of the senescence biomarker, p16^Ink4a^ (also known as Cdkn2a), impaired proliferation and resistance to apoptosis [6-10]. Senescent cells disrupt tissue structure and function because of the components they secrete, which act on adjacent as well as distant cells, causing fibrosis, inflammation, and a possible carcinogenic response[10]. Indeed, senescent cells possess a senescence-associated secretory phenotype (SASP), consisting of pro-inflammatory cytokines, chemokines and ECM-degrading proteins, which have deleterious paracrine and systemic effects [10-13]. Remarkably, even a relatively low abundance of senescent cells (10-15% in aged primates) is sufficient to cause tissue dysfunction [14].

To test whether senescent cells are causally implicated in age-related dysfunction and whether their removal is beneficial, J. L. Kirkland, T. Tchkonia, D. Baker (Mayo), J. van Deursen (Mayo) and colleagues made use of the biomarker for senescence, p16^INK4a^, and an inducible “suicide” gene designed by P. Scherer *et al*. [15] to develop a novel transgene, INK-ATTAC, to permit inducible elimination of p16^INK4a^-positive senescent cells upon administration of a drug (AP20187) [9]. In these mice, eliminating a relatively small proportion (~30%) of senescent cells extends health span and prevents the development of multiple age-related morbidities in both progeroid and normal, chronologically aged mice [9, 16-20]. Moreover, late life clearance attenuated the progression of already established age-related disorders [20]. In an effort to be applicable to humans, Kirkland and collaborators have identified a new class of drugs named senolytics. Through exploiting senescent cells’ dependence on specific pro-survival pathways, senolytics specifically kill senescent cells without affecting proliferating or quiescent, differentiated cells [21-23]. Recent studies have documented the use of senolytic drugs for the selective clearance of senescent cells from ‘aged’ tissues [16, 18-27]. Indeed, a combination of senolytics drugs (D, dasatinib, a FDA-approved tyrosine kinase inhibitor; and Q, quercetin, a flavonoid present in many fruits and vegetables), administered to 20-month-old male C57BL/6 mice once monthly for 4 months, led to significantly lower senescent (p16^Ink4a^- and SA-β-gal-expressing) cells in aorta, bone, adipose tissue, skeletal muscle, and kidney [11, 16-20, 24]. Whether eliminated genetically or through senolysis, it’s been shown that removal of p16^Ink4a^ senescent cells can delay the acquisition of age-related pathologies in adipose tissue, skeletal muscle, heart, blood vessels, lung, liver, bone, and eye [9, 11, 12, 16-20, 24-28]. Recently, the Kirkland lab has demonstrated that transplanting relatively small numbers of senescent preadipocyte cells into young (6 month old) mice causes persistent physical dysfunction, measured through maximal speed, hanging endurance and grip strength, 1 month after transplantation. Transplanting even fewer senescent cells into older (17 month old) recipients had the same effect and reduced survival, indicating the potency of senescent cells in shortening health- and lifespan. Intermittent oral administration of the senolytics, D and Q to senescent cell-transplanted young mice and naturally aged mice alleviated physical dysfunction and increased post-treatment survival by 36% while reducing mortality hazard to 65% [20]. Altogether these data indicate that cellular senescence is causally implicated in generating age-related phenotypes and that systemic removal of senescent cells can prevent or delay tissue dysfunction, physical dysfunction and extend health- and lifespan.

Mammalian aging is associated with gradual loss of the capacity of the tissue-specific stem/progenitor cells to maintain tissue homeostasis or to repair and regenerate tissues after injury or stress [29]. Indeed, in most tissues there is an overlap between aging and stem cell impairment [30-33]. Function of tissue-specific stem cells declines with age due to several factors including telomere shortening, increased senescence and elevated expression of p16^Ink4a^ and other cyclin-dependent kinase inhibitors (CDKIs) [34], DNA damage and external influences affecting stem cell niche homeostasis [30, 35-38]. Like other organs, the adult mammalian heart has the capacity, albeit low, to self-renew cardiomyocytes over the human lifespan [39], and genetic fate mapping models show that one source of new cardiomyocytes are a population of resident stem cells [40, 41]. Multiple labs [40, 42-56] have shown the mammalian heart, including human, to harbour a rare population of self-renewing, clonogenic, multipotent cardiac stem/progenitor cells (CSCs or CPCs; abbreviated hereafter as CPCs) with regenerative potential in vivo [40, 43-46, 48, 50, 56-58]. An undisputed mechanism of action of CPC transplantation is their secretome reparative potential, which through acting in a paracrine manner improves cardiomyocyte survival [59-61], limits fibrosis [60-62], induces angiogenesis [59, 61, 63], and new cardiomyocyte formation [62-64] and restores cardiac function [57, 59, 60, 64-66]. Recently, considerable confusion has emanated over the significant cardiomyogenic potential of CPCs purported using Kitcre Knockin or dual recombinases-mediated cell tracking mice models [67-69]. However, these models do not tag or specifically lineage trace the CPCs, neither have the investigators isolated, characterised or transplanted CPCs to test their stem cell and regenerative properties. Indeed, a recent communication showed that the very low number of endogenous c-Kit^pos^ CPC-generated cardiomyocytes detected in the Kitcre mice simply reflects the failure to recombine the CPCs to track their progeny and the severe defect in CPC myogenesis produced by the Kitcre allele [70].

Over the years, accumulated evidence from human and mouse studies demonstrate that cardiac aging and pathology affects the activity and potency CPCs [71-74]. This translates into a diminished capacity of the aged and diseased myocardium to maintain homeostasis, and repair and regenerate following injury [75-80]. The aging *milieu* might therefore limit the success of cell transplantation therapies where the outcome is direct cardiogenic differentiation of transplanted cells and/or stimulation of endogenous regenerative mechanisms. As the majority of cardiovascular disease patients in need of regenerative therapies are of advanced age, regulation of CPC and cardiovascular aging/senescence is mission critical.

Here we provide new information about the existence and biology of CPCs and further show their importance and relevance in the aged and diseased human heart, which we hope will help resolve the controversy plaguing this important field. Indeed, we have carried out an extensive analysis of CPCs in the human failing heart with advanced age and show the accumulation of senescent-CPCs, which exhibit diminished self-renewal, differentiation, and regenerative potential *in vivo*. We show that Senescent-CPCs have a SASP that negatively affects healthy non-senescent, cycling-competent CPCs, rendering them senescent. Clearing the senescent-CPCs using combinations of senolytic drugs attenuates the SASP and its effect on promoting senescence *in vitro*. The effects of global elimination of senescent cells on the heart and its regenerative capacity have not been elucidated. We report novel data that show systemic elimination of senescent cells *in vivo* in aged mice using senolytics (D+Q) or using the ‘suicide’ transgene, INK-ATTAC with administration of AP20187, results in CPC activation and increased number of small, immature, proliferating cardiomyocytes in the aged mouse heart.

## Results

### CPCs exhibit a senescent phenotype with increased age

Human CPCs were isolated from biopsies of right atria, obtained from subjects who had given informed consent before undergoing cardiac surgery (aortic disease, valve disease, coronary artery bypass graft (CABG), or multiple diseases), using sequential enzymatic digestion and dissociation, Optiprep density gradient to remove large debris, followed by magnetic activated cell sorting (MACS) (**Supplementary Figure 1a**). CPCs were magnetically enriched based upon a CD45-negative, CD31-negative, CD34-negative, and c-kit-positive sorting strategy [81] (**Supplementary Figure 1b**). Despite being recognised as a CPC marker, cells were not sorted for Sca-1 because its homology has not been confirmed in any species other than mouse. By flow cytometry analysis, CPCs showed expression of other recognised CPC markers such as CD90 (37±0.4%), CD166 (41±1%), CD105 (13±1%), and CD140α (5±0.4%) (**Supplementary Figure 1c**).

We isolated CPCs from 35 subjects of different genders, ages, and pathologies and found a linear increase (R^2^=0.722) in the number of CPCs that expressed the senescence-associated marker, p16^INK4A^, with age (**Figure 1a**). No differences were evident between males and females for p16^INK4A^ expression or between aortic disease, valvular disease, coronary artery disease, and multiple other diseases for p16^INK4A^ expression (**Supplementary Figure 2a,b**). On average, 22±9%, 31±4%, 48±9%, and 56±16% of CPCs expressed p16^Ink4a^ isolated from 50-59, 60-69, 70-79, and 80-89 year old subjects, respectively. We also found an increase (P<0.05) in the number of senescence-associated β-galactosidase- (SA-β-gal; ~60%) and DNA damage marker, γH2AX-positive CPCs (~20%) isolated from old (71-79 years), compared to middle-aged (54-63 years) subjects (**Figure 1b,c**). Moreover, p16^INK4A^-positive CPCs co-expressed γH2AX (**Figure 1c**). Further interrogation by Q-FISH revealed that, while the average telomere length of CPCs isolated from old and middle-aged subjects’ hearts were comparable, CPCs isolated from old (78-84 years) subjects’ hearts contained a 12% subpopulation with telomere length of <6Kb, which is regarded as being critically short (**Figure 1d**) [82]. Approximately 2% of the CPCs isolated from human hearts were Ki67-positive, reflective of their mainly dormant, quiescent phenotype [83]. There were no differences between middle-aged and old subjects in number of Ki67-positive CPCs, and we did not see any Ki67-positive CPCs that were p16^INK4A^-positive (**Supplementary Figure 2c**). These findings indicate that the aged human heart contains an increased proportion of aged senescent-CPCs, which could translate to their dysfunctionality.

**Figure. 1.**
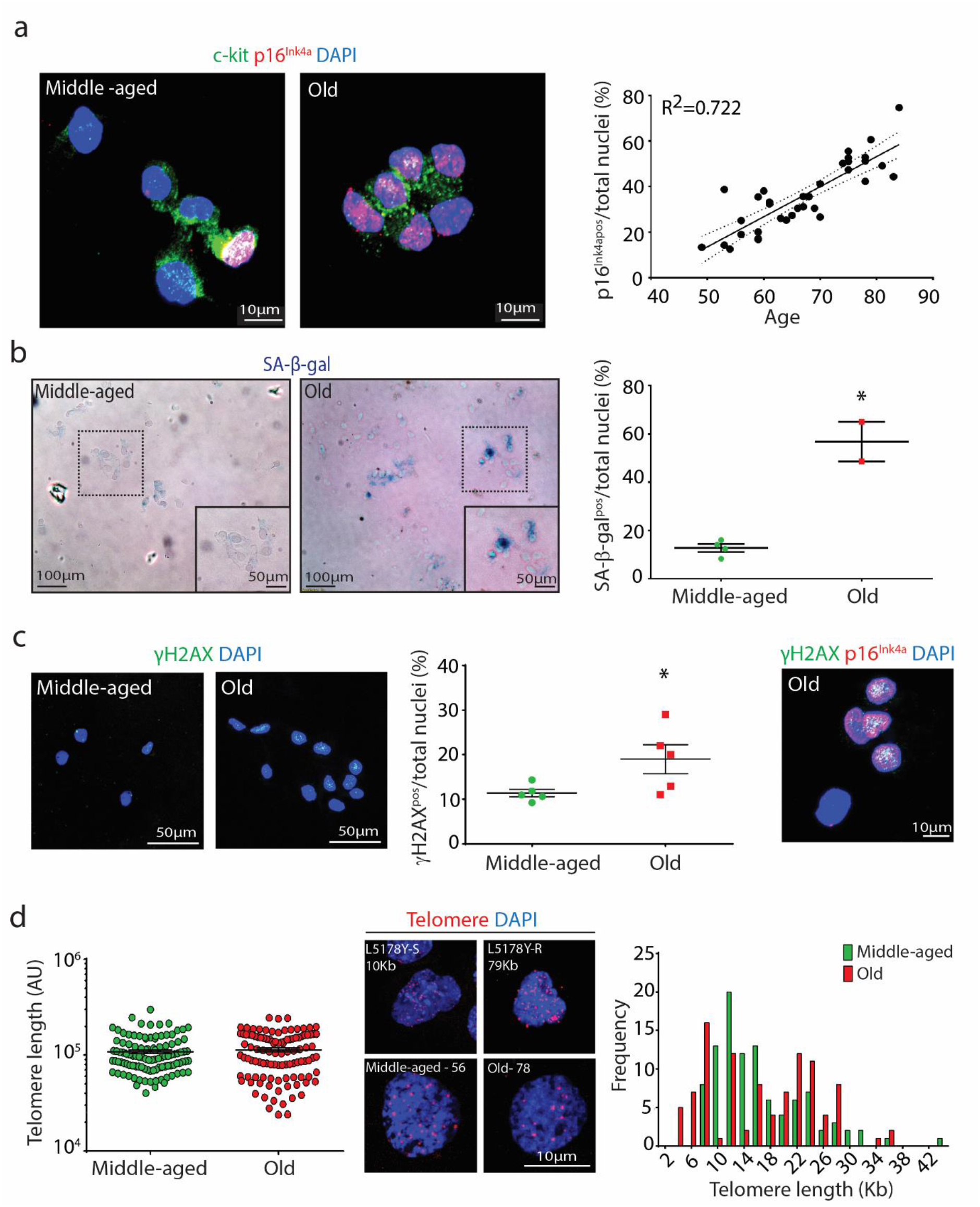
Over half of CPCs in the aged human heart are senescent. (**a**) Representative immunofluorescence images and quantification of freshly isolated c-kit^pos^ CPCs (green), p16^INK4A^-positive (red) expression, each data point represents the mean number of p16^INK4A^ CPCs per total nuclei for each individual donor (R^2^=0.722, n=32). (**b**) Representative immunofluorescence images and quantification of freshly isolated c-kit^pos^ CPCs stained for SA-β-gal (blue), each data point represents the mean number of SA-β-gal^pos^ cells per total nuclei for each individual donor (\**P*=0.0014, n=4 middle-aged, n=2 old). (**c**) Representative immunofluorescence images and quantification of freshly isolated c-kit^pos^ CPCs stained for γ-H2AX (green), each data point represents the mean percentage of γ-H2AX^pos^ cells per total nuclei for each individual donor (\**P*=0.0264, n=5). Nuclei stained in blue by DAPI. (**d**) Representative immunofluorescence images and quantification of freshly isolated c-kit^pos^ CPC Q-FISH telomere staining, each data point represents the telomere signal intensity (red) per total nuclei (DAPI, blue) calculated from 20 cells for each individual donor (n=5). Mouse cell lines with known telomere length, L5178Y-S (10 Kb) and L5178Y-R (79 Kb), were used to determine telomere length histograms (n=5). In each scatter graph, data points represent individual measurements and error bars represent SEM. Statistical analysis by Student’s t-test.

### CPCs from old subjects show impaired cell growth and differentiation

CPCs were isolated from 5 old (76-86 years) and 8 middle-aged (32-66 years) subjects, plated in growth medium, and propagated, where possible, to passage 11. CPCs isolated from 2 of the middle-aged (32 and 61 years) and 3 of the oldest (78, 80, and 86 years) subjects failed to grow and become established *in vitro*. Of the CPC cultures that did grow from all age groups (n=8), the CPCs, from P3 to P11, gradually lost their p16^INK4A^–positive subpopulation (**Supplementary Figure 2d,e**), likely due to the cell culture activated, cycling-competent CPCs outgrowing their p16^INK4A^–positive senescent counterparts. CPCs maintained the phenotype of c-kit-positive, CD31-negative over culture passage (**Supplementary Figure 2f**). To ensure that the effect of donor age could be effectively evaluated, all *in vitro* cell dynamic assays were performed between P2-P4.

CPCs isolated from old (77-86 years) subjects showed decreased (P<0.05) proliferation compared to CPCs isolated from middle-aged (34-62 years) subjects (**Figure 2a**). CPCs deposited as a single-cell in a 96-well plate generated a greater number (P<0.05) of clones if the single CPCs originated from middle-aged (34-62 years) subjects, compared to old (76-86 years) subjects (**Figure 2b**). Likewise, CPCs deposited at low dilution in bacteriological dishes for the generation of spheres in suspension were greater in number and size (P<0.05) for middle-aged (34-51 years) subject’s CPCs, compared to old (76-77 years) subject’s CPCs (**Figure 2c,d**). When CPCs were plated in cardiomyocyte differentiation medium they primed towards a cardiomyocyte-like precursor cell type that was Nkx2.5-positive, sarcomeric actin-positive, and Middle-aged (47-62 years) subject’s CPCs had increased (P<0.05) differentiation potential, compared to CPCs from old (76-77 years) subjects (**Figure 2e-g**). Older subject’s differentiated CPC-derived precursor cells showed disorganised sarcomeric structure (**Figure 2e**) and decreased (P<0.05) expression (**Figure 2g**), compared to differentiated CPCs from younger, middle-aged subjects.

**Figure. 2.**
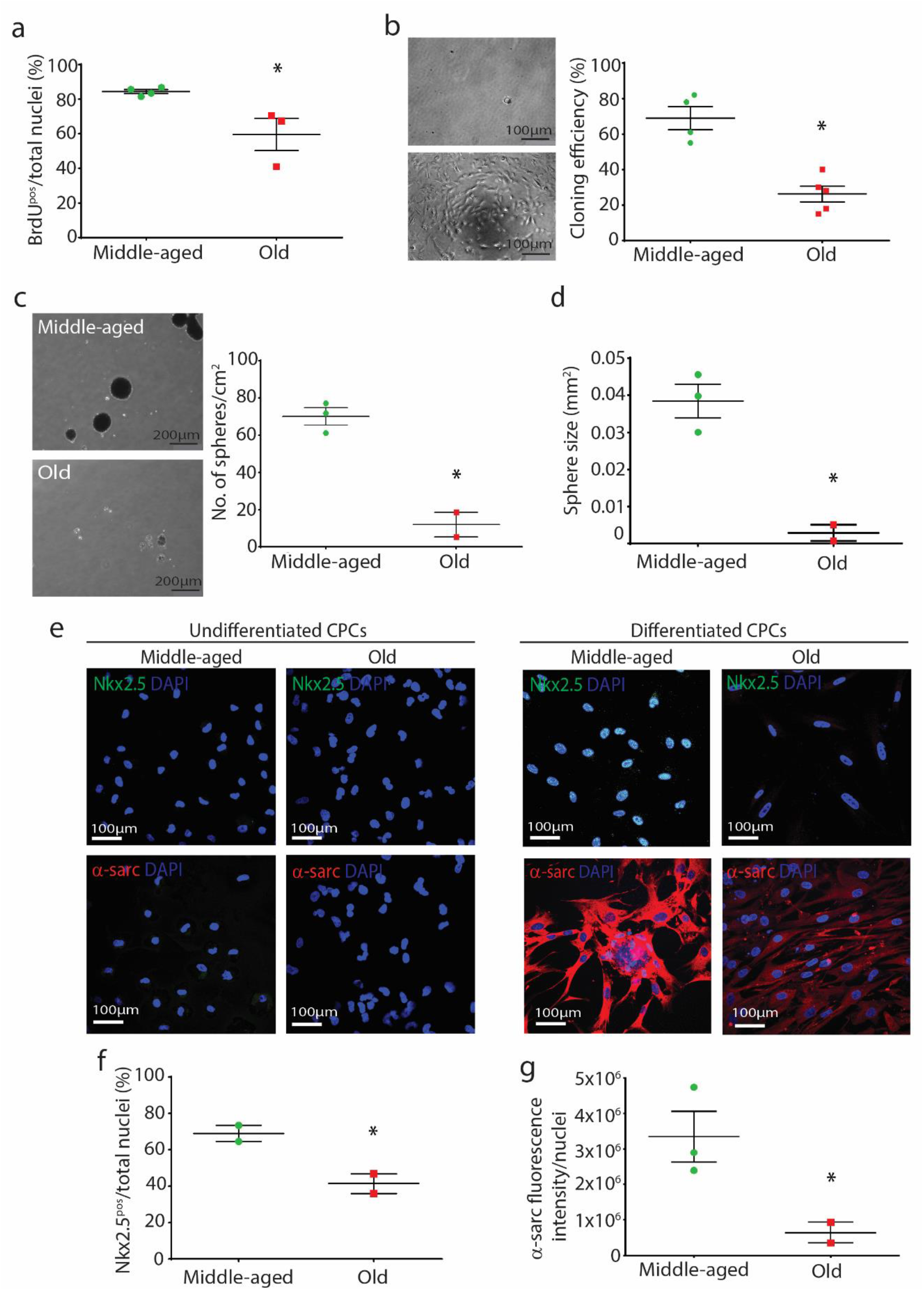
CPCs isolated from aged hearts exhibit diminished proliferation, clonogenicity, and cardiomyocyte differentiation potential. (**a**) CPC proliferation was determined using a BrdU incorporation assay, each data point represents the mean number of BrdU^pos^ cells per total nuclei for each individual donor (\**P*=0.0266, n=4 middle-aged, n=3 old). (**b**) Representative micrographs and quantification of a single CPC (top) and CPC-derived clone (bottom), each data point represents the mean cloning efficiency for each individual donor (\**P*=0.0008, n=4 middle-aged, n=5 old). (**c**) Representative micrograph and quantification of CPC-derived spheres from middle-aged (top) and old CPCs (bottom), each data point represents the mean number of spheres for each individual donor (\**P*=0.0051). (**d**) Quantification of the mean sphere size per donor (\**P*=0.01, n=3 middle-aged, n=2 old). (**e**) Representative immunofluorescence images of old and middle-aged donors’ CPCs stained αsarcomeric actin (red), Nkx2.5 (green) after 14 days of differentiation. Nuclei stained in blue by DAPI. (**f**) Quantification of Nkx2.5^pos^ cells per total nuclei for each individual donor (\**P*=0.0295, n=2). (**g**) Quantification of α-sarcomeric actin fluorescence intensity per nuclei for each individual donor (\**P*=0.023, n=3 middle-aged, n=2 old). In each scatter graph, data points represent individual measurements and error bars represent SEM. Statistical analysis by Student’s t-test.

Even though CPCs isolated from old hearts showed decreased proliferation, clonogenicity, and differentiation potential, only ~50% of CPCs are senescent in old myocardium (**Figure 1a**), therefore these data imply that a functionally cycling-competent CPC population still exists in old myocardium. Indeed, single CPC-derived clones from young, middle-aged, and old subjects were indistinguishable in terms of morphology, senescence, multipotency, self-renewing transcript profile, and differentiation (**Supplementary Figure 3**). These findings suggest that CPCs age and become senescent in a stochastic, non-autonomous manner. This resembles what was seen in rat preadipocytes [84].

### Aged-senescent CPCs lose their regenerative capacity *in vivo*

To purify for a senescent population of CPCs we utilised the C_12_-5-Dodecanoylaminofluorescein Di-*β*-D-Galactopyranoside (C12FDG) probe and pulled out SA-β-gal-positive CPCs through FACS (**Supplementary Figure 4a-d**). We also induced senescence in CPCs pharmacologically using Doxorubicin and Rosiglitazone (**Supplementary Figure 4e-g**), which we have used previously to render cells to senescence *in vitro* [11, 20]. Senescent-CPCs, whether Doxorubicin-, or Rosiglitazone-induced or purified using the C12FDG probe, exhibited a senescent phenotype being p16^I^NK4A–positive, Ki67-negative, and with shorter telomeres (**Supplementary Figure 5**). Senescent SA-β-gal-positive CPCs were non-proliferative and did not form clonal colonies when deposited as single cells, or generate spheres *in vitro*, compared to SA-β-gal-negative, Ki67-positive cycling-competent CPCs (**Supplementary Figure 5**). FACS phenotyping revealed decreased surface expression of the progenitor markers, c-kit, CD90, CD105, and CD166 and increased expression of CD31 and CD34 in SA-β-gal-positive, senescent-CPCs compared to SA-β-gal-negative, cycling-competent CPCs (**Supplementary Figure 6**).

To determine whether the dysfunctional stem/progenitor cell properties of the senescent-CPCs translated *in vivo*, we tested the regenerative capacity of senescent SA-β-gal-positive CPCs and cycling-competent SA-β-gal-negative CPCs in the myocardial infarction-regeneration mouse model (**Figure 3a**). Male, immunodeficient NSG mice were subjected to permanent ligation of the left anterior descending (LAD) coronary artery. Immediately after ligation, 5 x 10^5^ SA-β-gal-positive senescent or SA-β-gal-negative cycling-competent CPCs were injected intramyocardial in 15µl of PBS at 2 sites in the border zone. To serve as a cell control, a separate set of MI-mice were injected with 5 x 10^5^ non-CPCs (c-kit^neg^ cardiac-derived cells; containing 86±5% cardiac fibroblasts, 13±3% vascular smooth muscle, 1±1% endothelial cells [40]). Sham animals were treated the same way, except ligation of LAD coronary artery was not performed and they did not receive cells but were injected with the same volume of PBS. Mice were administered BrdU *via* osmotic mini pumps for 14 days after MI and cell injection to track new cell formation (**Figure 3a**). All cell populations were labelled prior to injection with PKH26 lipophilic membrane dye, which exhibited high labelling efficiency and label dye retention over population doublings in cycling-competent CPCs and c-kit^neg^ cardiac-derived cells *in vitro* (**Supplementary Figure 7**). We sacrificed a sub-set of MI-mice that had been injected with 5 x 10^5^ SA-β-gal-negative cycling-competent CPCs at 4 days. There was high engraftment and survival of CPCs within the infarct/border zone at 4 days (**Figure 3b**) and by 28 days the engraftment was still ~10% of cycling-competent CPCs per total nuclei in the infarct/border zone (**Figure 3c**). The engraftment and survival of senescent-CPCs and c-kit^neg^ cells at 28 days was significantly (P<0.05) less (**Figure 3c**).

**Figure. 3.**
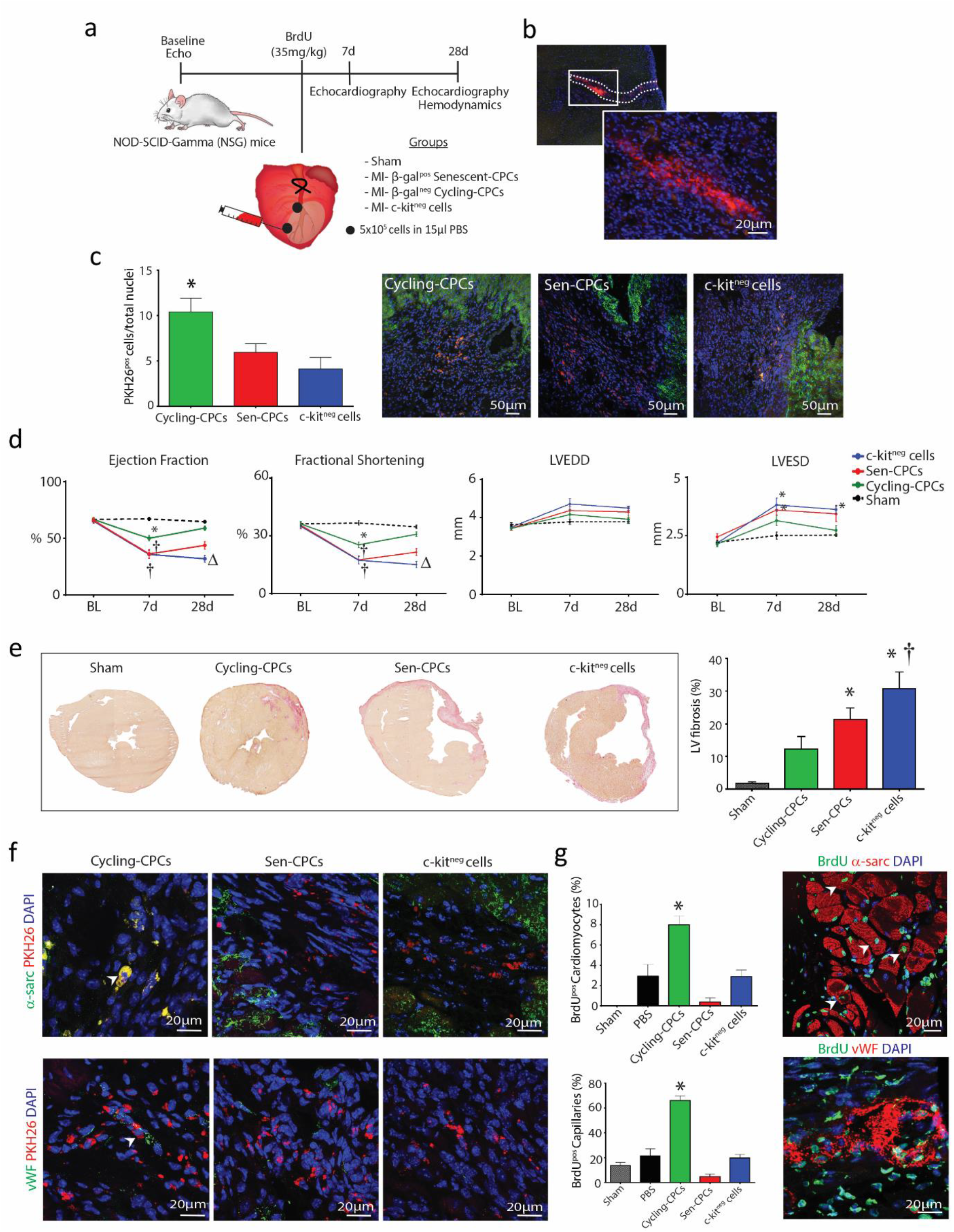
Aged-senescent CPCs show decreased reparative potential. (**a**) Experimental design for Nod-Scid-Gamma (NSG) mice left anterior descending (LAD) coronary artery ligation and treatment regime. (**b**) Representative immunohistological images showing PKH26 labelled CPCs in the myocardium 4 days post-MI. (**c**) Representative immunohistological images and quantification of engraftment rate of PKH26^pos^ cells (red) within the myocardium (α-sarcomeric actin; green) 28 days post-MI (\**P*=0.015 *vs*. c-kit^neg^ cells, n=4-5). (**d**) Echocardiography measurements of NSG mice at 7 and 28 days after MI and cell injection, compared to baseline (BL) showing LV ejection fraction (EF), fractional shortening (FS), left ventricular end-diastolic diameter (LVEDD), and left ventricular endsystolic diameter (LVESD) for Sham (black), MI-Cycling-competent CPCs (green), MI-Sen CPCs (red) and MI-c-kit^neg^ cells (blue) (\**P*<0.05 *vs*. Sham, †*P*<0.05 *vs*. MI-cycling-competent CPCs, Δ*P*<0.05 *vs*. MI-Sen CPCs, n=5-7). (**e**) Representative micrographs and quantification of HVG fibrosis staining in treated hearts (\**P*=0.0105 Sen CPCs *vs*. Sham, (\**P*=0.0003 c-kit^neg^ cells *vs*. Sham, †*P*=0.0112 *vs*. MI-cycling-competent CPCs, n=5-6). (**f**) Representative immunohistological images showing PKH26^pos^ cycling-competent CPCs co-expression of α-sarcomeric actinin (green) and vWF (green) (arrowheads) 28 days post-MI. (**g**) Representative immunohistological images and quantification of newly formed BrdU^pos^ (green) α-sarcomeric actinin^pos^ cardiomyocytes (red) and BrdU^pos^ (green) vWF^pos^ capillaries (red) in treated hearts (\**P*<0.05 *vs*. all treatment groups, n=5-7). Nuclei stained in blue by DAPI. All data are means ± SEM. Statistical analysis by one-way ANOVA with Tukey’s multiple comparison test.

At 1 week after LAD ligation all groups had decreased (P<0.05) LV function, compared to baseline and sham controls, however the group that had been injected with cycling-competent CPCs had less of a decrease in LV function at 1 week, compared to the senescent-CPC and non-CPC c-kit^neg^ cell groups (**Figure 3d**). The degree of MI represented as the % Average Area at Risk (AAR) through Evans Blue staining immediately after MI was 33.0±1.6% (n=5), demonstrating the operator as being extremely consistent in inducing a similar size MI injury to each mouse. At 4 weeks after LAD ligation, MI hearts that had received cycling-competent CPCs showed an improvement (P<0.05) in LV ejection fraction (EF), fractional shortening (FS), LVEDD, and LVESD, which had almost returned to baseline values and to that of Sham controls (**Figure 3d**).

This extent of LV improvement was not apparent in MI-mice that were injected with SA-β-gal-positive senescent-CPCs, or the non-CPC c-kit^neg^ cell group that showed no recovery with worsened LV function and were all in heart failure at 4 weeks (**Figure 3d**). To accompany these functional changes, cycling-competent CPC injection resulted in a decreased (P<0.05) infarct size, whereas SA-β-gal-positive senescent-CPCs or non-CPC c-kit^neg^ cells did not change the extent of infarct size (**Figure 3e**). Immunohistochemical analysis of cross-sections revealed that at 4 weeks after MI, the transplanted PKH26-labelled cycling-competent CPCs had increased expression of sarcomeric proteins, α-actinin, as well as the endothelial lineage marker vWF, evidencing their differentiation into cardiomyocyte-like precursor cells and endothelial cells, respectively (**Figure 3f**). Limited or no differentiation was evident in the infarcted/border zone of the hearts injected with PKH26-labelled senescent-CPCs and non-CPC c-kit^neg^ cells (**Figure 3f**). To determine whether the transplanted cells had participated in inducing a paracrine effect, the infarct/border zone of hearts that had received cells were analysed for formation of new cells that were BrdU-positive/PKH26-negative. Hearts that were injected with cycling-competent CPCs showed an increased number of BrdU-positive cells, compared to those injected with SA-β-gal-positive senescent-CPCs or non-CPC c-kit^neg^ cells (**Supplementary Figure 8**). Moreover, these BrdU-positive cells co-localised with vWF or α-sarcomeric actinin, indicating new endothelial (capillary) cell and cardiomyocyte formation, respectively. New cardiomyocyte and capillary formation was more evident (P<0.05) in the MI-cycling-competent CPC group (**Figure 3g**). These findings show the diminished regenerative and reparative capacity of senescent CPCs, compared to healthy, cycling-competent CPCs.

### Aged-Senescent CPCs have a Senescence-Associated Secretory Phenotype (SASP)

Senescent cells exhibit a SASP [10]. Senescent SA-β-gal-positive CPCs showed increased expression of SASP factors, including MMP-3, PAI1, IL-6, IL-8, IL-1β, and GM-CSF, compared to non-senescent, SA-β-gal-negative, cycling-competent CPCs (**Figure 4a**). To determine whether the SASP factors were secreted from senescent-CPCs, we quantified the protein levels of seven of the highly expressed SASP factors in conditioned media from senescent-CPCs using Luminex technology. We found increased (P<0.05) quantities of all seven SASP factors in senescent-CPC conditioned medium, compared to conditioned medium of cycling-competent CPCs (**Figure 4b**). Next we treated cycling-competent CPCs with conditioned medium from senescent-CPCs and measured cell proliferation and senescence of the cycling-competent CPCs. Conditioned medium from senescent-CPCs resulted in decreased (P<0.05) proliferation (**Figure 4c**) and an increased (P<0.05) proportion of senescent p16^INK4A^–positive, SA-β-gal-positive, and γH2AX-positive CPCs in the culture, compared to CPCs treated with conditioned medium from cycling-competent CPCs or unconditioned medium (**Figure 4d-f**). These findings show that the senescent-CPCs exhibit a SASP, which can negatively impact surrounding cells, rendering otherwise healthy, cycling-competent CPCs to lose proliferative capacity and switch to a senescent phenotype.

**Figure. 4.**
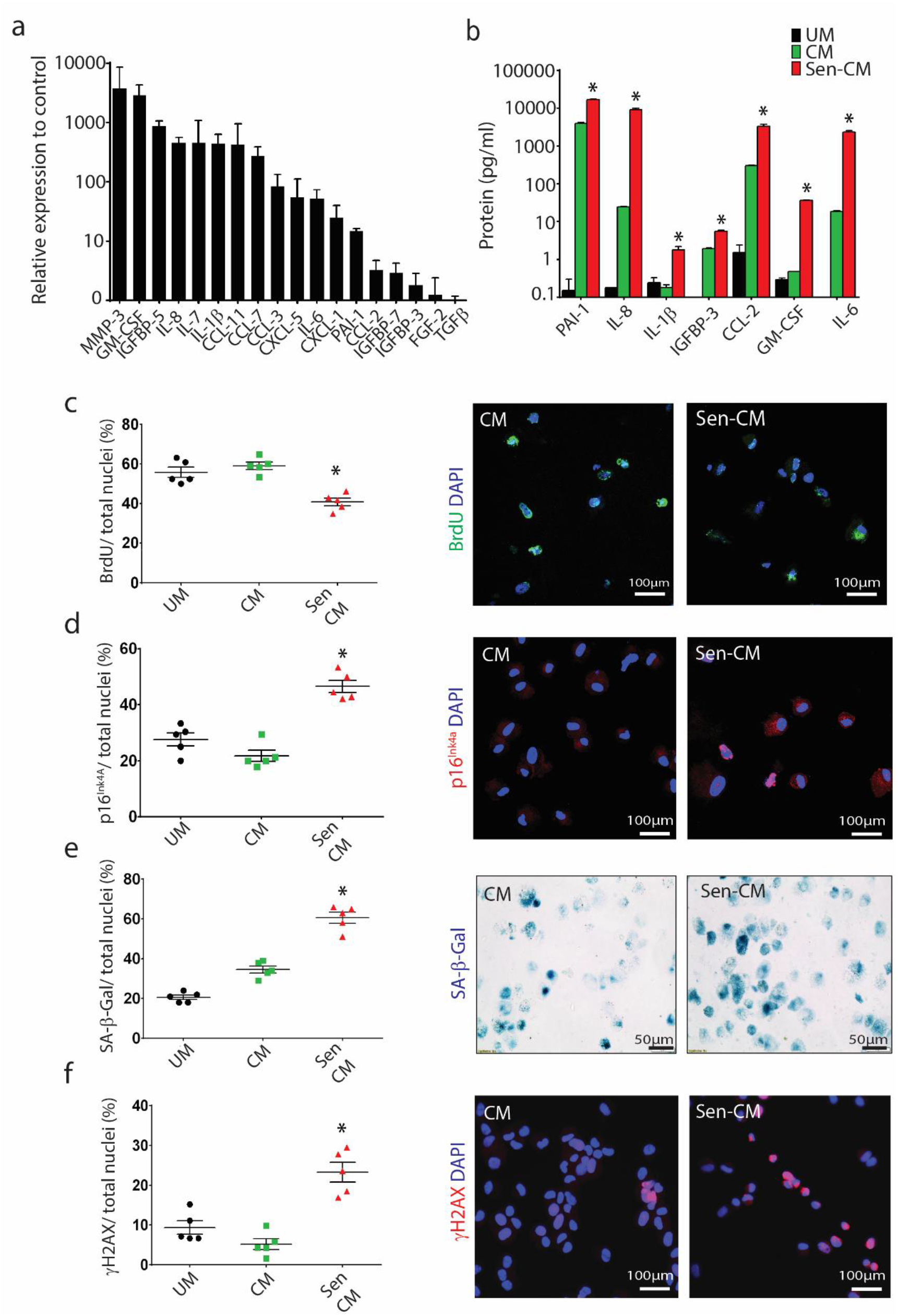
Aged-senescent CPCs have a SASP. (**a**) Gene expression of Senescent-CPC SASP factors relative to cycling-competent CPCs (control). (**b**) SASP factor protein levels quantified by Luminex analysis of unconditioned media (UM), Cycling-competent CPC (CM) and Senescent-CPC (Sen CM) conditioned media, PAI-1 (\**P*=0.0006), IL-8 (\**P*=0.003), IL-1ß (\**P*=0.0283), IGFBP-3 (\**P*=0.0011), CCL2 (\**P*=0.0116), GM-CSF (\**P*=0.0001), IL-6 (\**P*=0.0030) Sen CM *vs*. CM. Conditioned media applied to cycling-competent CPCs and the following analyses performed; (**c**) Quantification of BrdU^pos^ (green) CPCs per total nuclei and representative immunocytochemical images (\**P*=0.001 Sen CM *vs*. UM, \**P*=0.0002 Sen CM *vs*. CM). (**d**) p16^INK4A-pos^ (red) CPCs per total nuclei (\**P<*0.0001 Sen CM *vs*. UM, \**P*<0.0001 Sen CM *vs*. CM). (**e**) SA-β-gal^pos^ (blue) CPCs per total nuclei (\**P<*0.0001 Sen CM *vs*. UM, \**P*<0.0001 Sen CM *vs*. CM). (**f**) γH2AX^pos^ (red) CPCs per total nuclei (\**P*=0.0006 Sen CM *vs*. UM, \**P*<0.0001 Sen CM *vs*. CM). Nuclei stained in blue by DAPI. In each scatter graph, data points represent individual measurements and error bars represent SEM. Statistical analysis by one-way ANOVA with Tukey’s multiple comparison test.

### Elimination of senescent CPCs using senolytic drugs abrogates the SASP *in vitro*

Removal of p16^Ink4a^ senescent cells can delay the acquisition of age-related pathologies in adipose tissue, skeletal muscle, heart, blood vessels, lung, liver, bone, and eye [9, 11, 12, 16-20, 24, 25, 28]. Recent studies have documented the use of senolytic drugs for the selective clearance of senescent cells from ‘aged’ tissues [16-27]. We tested the potential of 4 senolytic drugs, Dasatinib (D; an FDA-approved tyrosine kinase inhibitor), Quercetin (Q; a flavonoid present in many fruits and vegetables), Fisetin (F; also a flavonoid), and Navitoclax (N; an inhibitor of several BCL-2 family proteins), alone and in combination to eliminate and clear senescent-CPCs *in vitro* (**Supplementary Figure 9a**). Measuring cell viability with crystal violet and number of SA-β-gal-positive CPCs, dose-response experiments on senescent- or cycling-competent CPCs from the same subjects showed D and N to effectively clear senescent-CPCs, whereas F and Q were less effective at clearing senescent-CPCs (**Supplementary Figure 9b,c**). However, D also decreased the viability of cycling-competent CPCs (**Supplementary Figure 9b**). A combination of D+Q, which has previously shown to yield effective senescent cell clearance [16-20, 24, 25] and which does not share the toxic anti-neutrophil and anti-platelet side effects of N [22, 23] was tested, and at a dose of 0.5µM D with 20µM Q, cycling-competent CPC viability was preserved (**Figure 5a**), but senescent-CPCs were cleared and induced to selective apoptosis (**Supplementary Figure 9d**).

**Figure. 5.**
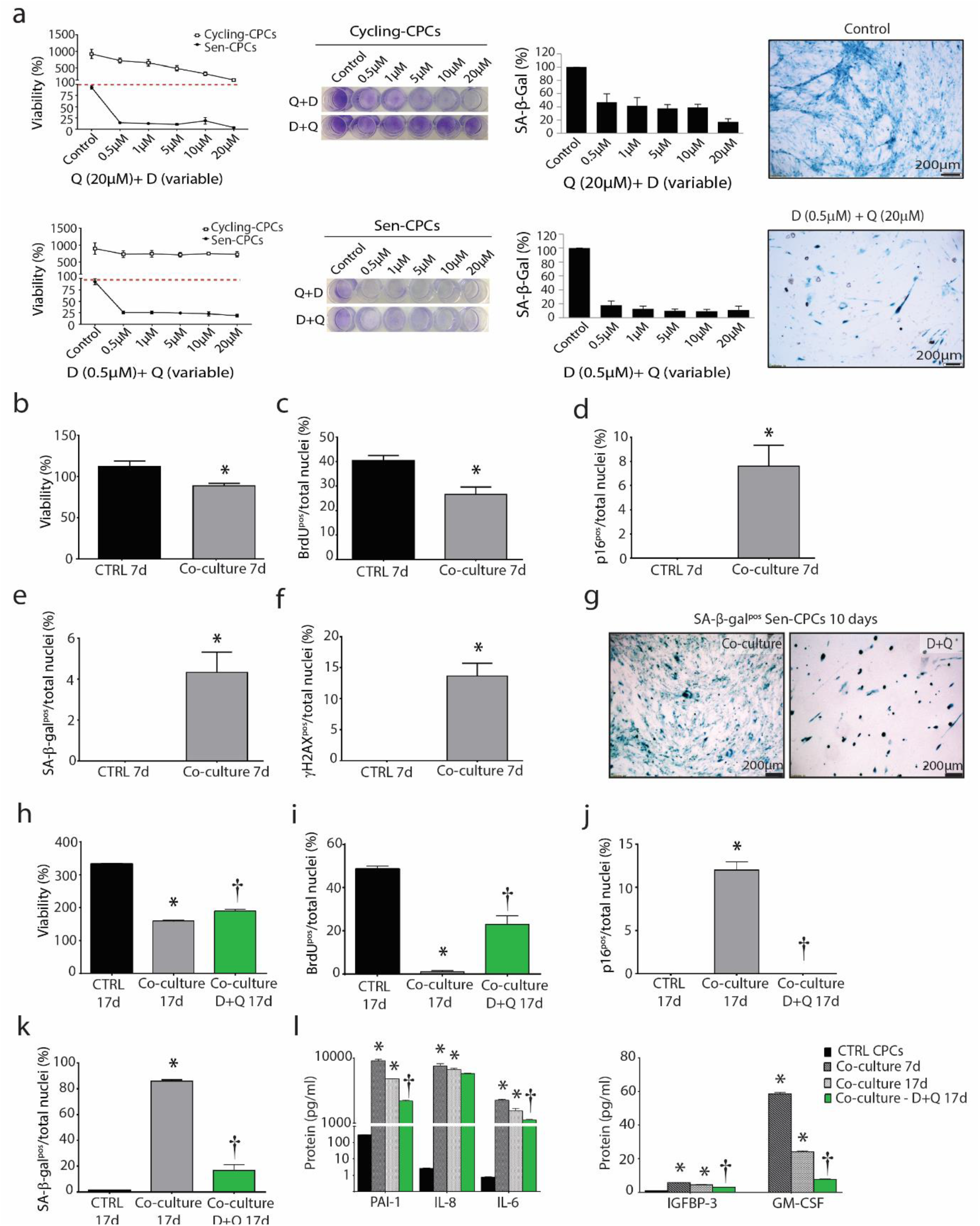
Senolytic clearance abrogates the SASP. (**a**) Quantification of % viability and % SA-β-gal staining of cycling-competent CPCs and Senescent-CPCs exposed to various concentrations of D+Q for 3 days. Representative crystal violet (cell viability) and SA-β-gal (cell senescence) staining. (**b-f**) Quantification of 7 days of co-culture of cycling-competent CPCs with Senescent-CPCs for (**b**) viability (\**P*=0.0068); (**c**) BrdU incorporation (\**P*=0.0056); (**d**) p16^INK4A^ (\**P*=0.0025); (**e**) SA-β-gal (\**P*=0.0025) and (**f**) γH2AX (\**P*=0.0002). CTRL is cycling-competent CPCs alone. Data are means ± SEM. Statistical analysis by Student’s t-test. (**g**) Representative SA-β-gal staining after clearance of Senescent-CPCs from co-culture by D+Q treatment. (**h-k**) Quantification of 17 days of co-culture of cycling-competent CPCs with Senescent-CPCs or co-culture with D+Q treatment for (**h**) viability (\**P*<0.0001 *vs*. CTRL 17d, †*P*=0.0001 *vs*. co-culture 17d); (**i**) BrdU incorporation (\**P*<0.0001 *vs*. CTRL 17d, †*P*<0.0001 *vs*. co-culture 17d); **(j**) p16^INK4A^ (\**P*<0.0001 *vs*. CTRL 17d, †*P*<0.0001 *vs*. co-culture 17d) and (**k**) SA-β-gal (\**P*<0.0001 *vs*. CTRL 17d, †*P*<0.0001 *vs*. co-culture 17d). CTRL is cycling-competent CPCs alone. Data are means ± SEM. Statistical analysis by a one-way ANOVA with Tukey’s multiple comparison test. (**l**) SASP factor protein levels quantified by Luminex analysis from each treatment condition; PAI-1 (*\*P*<0.0001 *vs*. CTRL, †*P*=0.0003 *vs*. co-culture 17d); IL-8 (*\*P*<0.0001 *vs*. CTRL); IL-6 (*\*P*<0.0001 *vs*. CTRL); IGFBP-3 (*\*P*<0.0001 *vs*. CTRL, †*P*<0.0001 *vs*. co-culture 17d); GM-CSF (*\*P*<0.0001 *vs*. CTRL, †*P*<0.0001 *vs*. co-culture 17d). Data are means ± SEM. Statistical analysis was performed by a one-way ANOVA.

We next determined whether clearing senescent-CPCs using D+Q would abrogate the SASP and its paracrine impact on CPCs. Using transwell inserts, cycling-competent CPCs were seeded on the top chamber insert and co-cultured in the presence of senescent-CPCs seeded on the bottom chamber. Cultures were left for 7 days and then cycling-competent CPCs in the top chamber were analysed for proliferation and markers of senescence, p16^INK4A^, SA-β-gal, and γH2AX, and conditioned medium analysed for SASP factors. The cultures were then treated with D+Q for 3 days to clear the senescent-CPCs on the bottom chamber, and then 7 days later cycling-competent CPCs in the top chamber were analysed for proliferation and the markers of senescence, p16^INK4A^, SA-β-gal, and γH2AX, and conditioned medium analysed for SASP factors (total of 17 days; **Supplementary Figure 9a**). We found that cycling-competent CPCs co-cultured in the presence of senescent-CPCs for 7 days were decreased (P<0.05) in number and proliferation, and had increased (P<0.05) expression of p16^INK4A^,SA-β-gal, and γH2AX (**Figure 5b-f**). Application of D+Q to co-cultures eliminated the senescent-CPCs (**Figure 5g**) and 7 days later, the cycling-competent CPCs had increased (P<0.05) in number (**Figure 5h**), proliferation (**Figure 5i**), and the number of p16^INK4A^ and SA-β-gal CPCs had decreased (P<0.05) compared to CPCs that had been in co-culture with senescent-CPCs for 17 days (**Figure 5j,k**). Co-culture of cycling-competent CPCs with senescent-CPCs led to increased (P<0.05) secretion of SASP factors into the medium, but the level of SASP factors was reduced (P<0.05) with application of D+Q (**Figure 5l**). These findings document that senescent CPCs have a SASP, and clearance of senescent CPCs using a combination of D+Q senolytics abrogates the SASP and its detrimental senescence-inducing effect on healthy, cycling-competent CPCs.

### Elimination of senescent cells *in vivo* activates resident CPCs and increases number of small, proliferating cardiomyocytes in the aged heart

Eliminating a relatively small proportion (~30%) of senescent cells using a ‘suicide’ transgene, INK-ATTAC, that permits inducible elimination of p16^Ink4a^-expressing senescent cells upon the administration of a drug (AP20187) extends health span and prevents the development of multiple age-related morbidities in both progeroid and normal, chronologically aged mice [9, 16-20, 28]. Moreover, in 20-month-old male C57BL/6 mice either randomized to vehicle or D+Q treatment once monthly for 4 months, D+Q treatment led to significantly lower senescent (p16^Ink4a^- and SA-β-gal-expressing) cells in bone, adipose tissue, skeletal muscle, and kidney [16-20, 24]. Here 24-32 month INK-ATTAC transgenic or wild-type mice were randomised to either vehicle, AP20187, or D+Q treatment, administered in 4 cycles for 3 consecutive days/cycle, 12 days apart. Mice were sacrificed 4 days after the last dose of cycle 4 (**Figure 6a**). Tissues in which p16^Ink4a^ expression and/or senescent cells are decreased by D+Q in wild type mice as well as by AP20187 in INK-ATTAC mice include the aorta, adipose tissue, cardiac and skeletal muscle, lung, liver, and bone [9, 16-28]. We showed p16^Ink4a^ mRNA expression was decreased (P<0.05) in the heart following D+Q or AP20187 treatment in aged INK-ATTAC or wildtype mice (**Figure 6b**).

Previously we have shown an improvement of heart function in old mice after D+Q treatment [24]. Analysis of cardiac cross-sections revealed significantly higher (P<0.05) CPC (Sca-1^+^/c-kit^+^/CD45^-^/CD31^-^/CD34^-^) [83] numbers (**Figure 6c; Supplementary Figure 10a**) in AP20187-treated INK-ATTAC mice and D+Q-treated INK-ATTAC or wild-type mice, compared to vehicle-treated control. Interestingly, D+Q treatment showed increased (P<0.05) CPC number, compared to AP20187-treatment (**Figure 6c**). Roughly 10% of CPCs were activated and in the cell cycle (Ki67-positive) at the time of sacrifice, after AP20187 or D+Q treatment (**Supplementary Figure 10b**). Morphometric analysis of heart sections showed that AP20187-treated and D+Q-treated mice had increased number of smaller ventricular myocytes (**Figure 6d**), suggesting these myocytes to be immature and newly formed, compared to vehicle-treated mice, which exhibited only rare small myocytes but a greater proportion of hypertrophied myocytes (**Figure 6d**). We found an increase (P<0.05) of small, proliferating Ki67-positive myocytes (~0.25%) in old hearts following AP20187- or D+Q-treatment, compared to vehicle-treated control (0.03±0.03%) (**Figure 6e,f**). To corroborate these data we injected EdU 4 days and 2 hours prior to sacrifice of old (22m) and young (3m) AP20187-treated INK-ATTAC mice (**Supplementary Figure 10c**). We found increased (P<0.05) number of small EdU-positive myocytes (0.25±0.06%; **Figure 6g,h**) in the hearts of old AP-treated mice, compared to old vehicle-treated (0.07±0.00%), young vehicle-treated (0.12±0.03%) and young AP-treated (0.11±0.04%) mice. The number of EdU-positive myocytes in the old AP-treated INK-ATTAC mice was the same in amount to Ki67-positive myocytes (~0.25%) in AP20187- or D+Q-treated mouse hearts (**Figure 6f**). Finally, we detected a decrease (P<0.05) in fibrosis in the LV following AP20187- and D+Q-treatment, compared to vehicle-treated control (**Figure 6i**). In contrast to the treatment of aged mice, treatment of young adult (2-3 Months) INK-ATTAC or wild-type mice with AP20187 or D+Q, respectively, did not alter EdU-positive myocyte number (**Figure 6h**), CPC numbers or myocyte diameter (*data not shown*). These findings show that clearance of senescent cells leads to stimulation of CPCs and cardiomyocyte proliferation and that this strategy is specific to the aged heart.

**Figure. 6.**
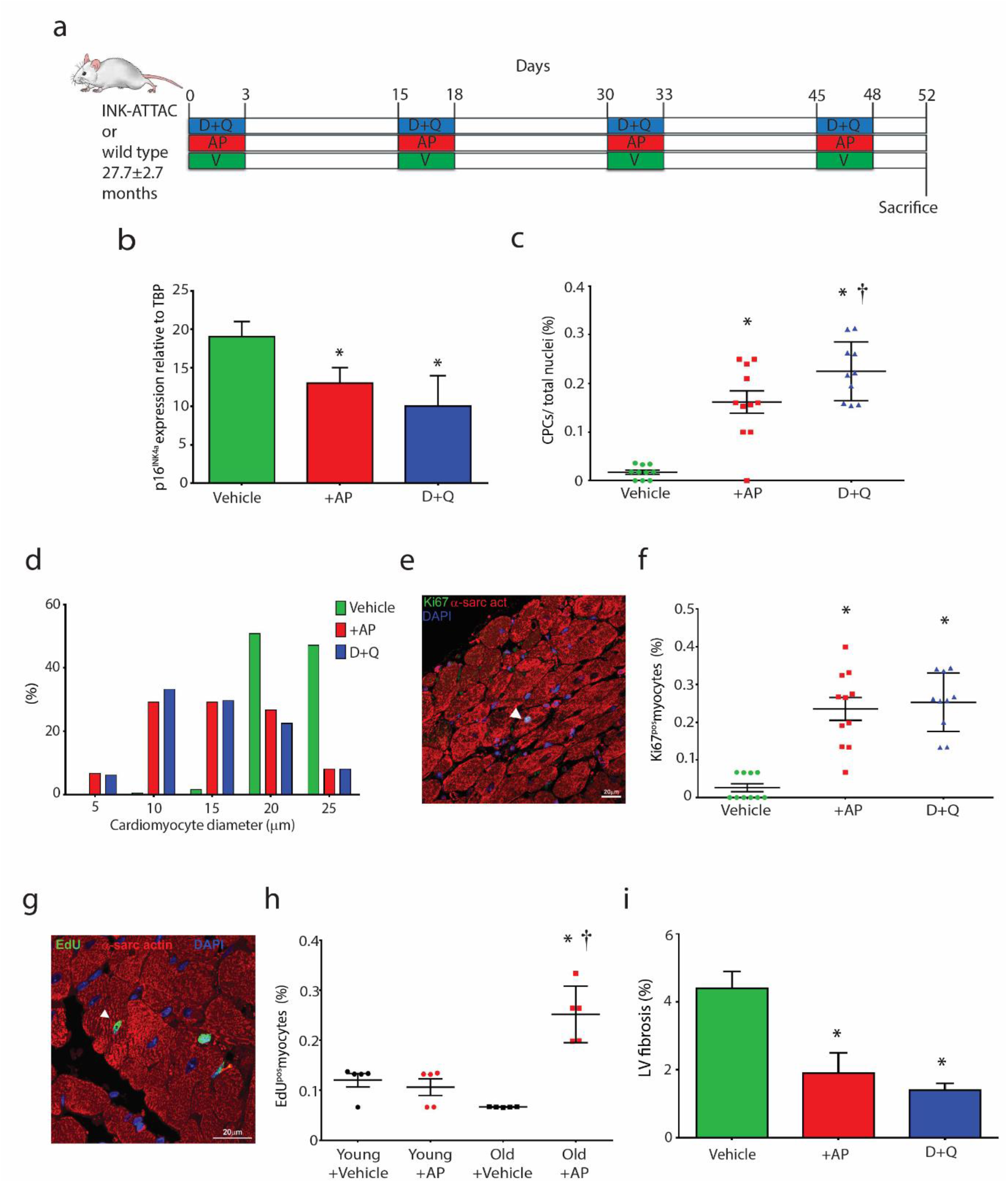
Clearance of senescent cells leads to stimulation of CPCs and new myocyte formation in the aged heart. (**a**) Experimental design for senescent cell clearance *in vivo* experiments. (**b**) Total p16^Ink4a^ gene expression relative to TBP in aged treated hearts (\**P*=0.0039, AP *vs*. Vehicle; *P=0.001, D+Q *vs*. Vehicle; n=5). (**c**) Quantification of CPCs (\**P*<0.0001 *vs*. Vehicle, †*P*=0.0453 vs. AP, n=10-11). Data points represent individual mice and error bars represent SD. (**d**) Frequency distribution histogram of cardiomyocyte diameter (n=6). (**e**) A Ki67-positive (green) cardiomyocyte (red) in the LV of a 32 month INKATTAC D+Q-treated mouse. Nuclei are stained by DAPI in blue. (**f**) Quantification of Ki67^pos^ myocytes per total myocytes (\**P*=<0.0001 AP *vs*. Vehicle, \**P*=<0.0001, D+Q *vs*. Vehicle, n=10). (**g**) An EdU-positive (green) cardiomyocyte (red) in the LV of a 22 month INK-ATTAC AP-treated mouse. Nuclei are stained by DAPI in blue. (**h**) Quantification of EdU^pos^ myocytes per total myocytes (\**P*==<0.0001 Old+AP *vs*. Old+Vehicle, †*P*=<0.0001 Old+AP *vs*. Young+AP, n=5). (**i**) Quantification of LV fibrosis (\**P*=0.0210 AP *vs*. Vehicle, \**P*=0.0092 D+Q vs. Vehicle, n=3). Data are Mean ± SD. Statistical analysis by one-way ANOVA with Tukey’s multiple comparison test.

## Discussion

In a large sample cohort our study shows that CPCs isolated from the failing human heart develop a senescent phenotype with age exhibited by increased expression of senescence-associated markers (p16^INK4A^, SA-β-gal), DNA damage, shortened telomere length and a SASP. Aged human hearts with dilated cardiomyopathy showed greater numbers of p16^INK4A^–positive CPCs and cardiomyocytes with shorter telomeres than age-matched controls [72]. Similarly, CPCs isolated from failing, aged hearts show increased p16^INK4A^ and inflammatory factor expression [73]. Reliably detecting senescent cells in vivo is an ongoing challenge, and it is important that a combination of senescent cell biomarkers are used for detection as any one marker used in isolation is prone to false positives. Our study used a combined panel of the senescence-associated biomarkers, p16^INK4A^, γH2AX, telomere length, SA β-gal activity and SASP expression, to detect senescent CPCs. Our findings demonstrate that CPCs accumulate in the failing hearts of elderly subjects (>76 years) and are dysfunctional, showing impaired proliferation, clonogenicity, spherogenesis, and differentiation, compared to CPCs isolated from the hearts of middle-aged (32-66 years) subjects. As the adult heart possesses very low numbers of cardiomyogenic CPCs [83], if by 80 years of age >50% of resident CPCs are senescent, this presents a bleak outcome for harvesting healthy, functional CPCs from patients who are candidates for regenerative therapies and their autologous use. Moreover, strategies to activate the regenerative capacity of the aged heart through delivery of growth factors or cell therapy will also be sub-optimal. Therefore, the success of cardiac regenerative therapeutic approaches thus far tested for treating patients with heart failure and disease could be of limited efficacy in promoting myocardial regeneration because of the increased number of senescent, dysfunctional CPCs and cardiomyocytes [72, 73] and the resultant presence of a cardiac SASP in the aged and failing heart that impairs the function of the remaining non-senescent CPCs.

The present study found that CPCs age in a stochastic non-autonomous manner and it is possible to clonally select for a cycling-competent population of CPCs even from diseased or aged hearts. There are individual CPCs in older individuals that have replicative and functional capacities resembling those of CPCs in younger subjects. A similar scenario was found in the case of rat fat cell progenitors [84]. While the abundance of progenitors cloned from adipose tissue that had restricted capacities for replication and differentiation into adipocytes or that were non-replicative but viable (*i.e*., senescent) increased progressively with aging in rat fat, there remained cells that had the capacities for replication and adipogenic differentiation characteristic of clones derived from young rats. Together, these findings indicate that: (1) it may be feasible to isolate CPCs even from older individuals that are functional, capable of supporting cardiac regeneration if removed from their toxic *milieu*, and that could be therapeutically relevant in treating patients, especially if they were autologously generated [85] and (2) that by clearing senescent CPCs with a toxic SASP from the aged heart, there remains a tissue-resident population of CPCs with capacity to regenerate damaged heart tissue.

When we purified for a homogenous SA-β-gal-positive, senescent CPC population, we showed that these cells had poor engraftment and survival, and were unable to contribute to cardiac regeneration, repair or restoration of cardiac function following transplantation into the infarcted myocardium. This is contrary to *in vitro*-selected, SA-β-gal-negative, proliferative cycling-competent (Ki67^pos^) CPCs, which had high survival and engraftment in the infarct/border zone, restored cardiac function almost to baseline and sham control values (LVEF 59±2% at 28 days vs. 66±2 at baseline), decreased infarct size, differentiated into endothelial and cardiomyocyte-like precursor cells as well as enhanced endogenous new cardiomyocyte and capillary formation. Injection of CPCs that show high survival and engraftment [60], and have the ability to generate new cardiomyocytes and vasculature [40, 48, 81, 83] can facilitate physiologically significant repair and regeneration, with resultant improvements in cardiac function in both humans [66] and rodent models [40, 43, 45, 48, 61, 83, 86]. Although some of the transplanted CPCs expressed α-sarcomeric actin, these cells did not exhibit the typical cardiomyocyte phenotype as they were small and lacked a structured sarcomeric unit. Therefore, they could not be considered as new, immature myocytes which contributed physiologically to the substantially improved LV function. There is now a general consensus that the favourable effect of cell transplantation protocols is, at least in part, mediated by ‘paracrine’ effectors secreted by the transplanted cells, contributing to improved myocardial contractility and amelioration of ventricular remodelling (decreasing fibrosis, hibernation, and stunning), inhibition of the inflammatory response, increased cardiomyocyte survival, cardiomyogenesis and angiogenesis/neovascularisation [87]. The present data emphasize the importance of taking into account the hostile infarcted environment, which does not favour engraftment, differentiation or maturation of newly formed cardiomyocytes derived from injected cells. However, the presence of CPC-derived cardiomyocyte precursors expressing sarcomeric protein in the infarct/border zone is promising and further work should elucidate how to mature these cells into functionally competent contractile cells.

Like the present study, not all experimental studies have shown physiological regeneration of CPC-derived cardiac muscle following administration of CPCs [60, 88]. This is most likely due to the heterogeneous nature of cardiac c-kit positive cells tested, with only a very small fraction (1-2%) being cardiomyogenic [83]. Bringing together the cardiomyogenic potential of CPCs, which can be amplified in number through *in vitro* clonal selection [40, 83] and CPC-mediated cytokine release that regulates cardiac cells’ behavior, advocate CPCs as an ideal cell candidate for cardiac regenerative therapies.

Senescent cells have emerged as bona fide drivers of aging and age-related CVD, which suggests strategies aimed at reducing or eliminating senescent cells could be a viable target to treat and prevent CVD [89]. Baker et al. (2016) showed in 18 month old INK-ATTAC mice that p16^INK4a^-positive cells contribute to cardiac aging and these senescent cells decreased following AP20187-treatment [28]. We show for the first time that the genetic and pharmacological approaches used here to reduce senescent cell burden leads to activation of the endogenous regenerative capacity in the aged heart. AP20187-treated INK-ATTAC and D+Q-treated INK-ATTAC and wild-type aged mice had an increased number of CPCs and smaller ventricular myocytes, which were Ki67-positive and EdU-positive, suggesting these myocytes to be immature and newly formed, compared to vehicle-treated mice, which exhibited very rare small myocytes but a greater proportion of hypertrophied myocytes. These findings are in line with those of Baker et al. (2016) who showed that AP-treated INK-ATTAC mice had smaller ventricular cardiomyocytes.

The frequency of resident cardiac stem/progenitor cells in the healthy myocardium of several mammalian species, including human, mouse, rat, and pig, is approximately one per every 1000–2000 myocytes, depending on age [90]. We detected a 16- and 23-fold increase in the number of CPCs following elimination of senescent cells by AP- or D+Q-treatment in the aged mouse heart, respectively. Moreover, ~10% of CPCs after elimination of senescent cells were activated, expressing Ki67. The number of Ki67-positive and EdU-positive cardiomyocytes increased 9- and 4-fold in the aged heart, respectively, following clearance of senescent cells by either D+Q- or AP-treatment. The number of proliferating cardiomyocytes present in the young (3 month old) mouse heart is 0.12±0.03% of total cardiomyocytes, and the number of proliferating cardiomyocytes present in the old (22 month old) mouse heart is 0.07±0.00% of total cardiomyocytes. Elimination of senescent cells lead to double the amount of the proliferating cardiomyocytes found in a young heart, and triple the number found in an old heart. Therefore, the present data represent a significant and physiologically relevant increase and activation of the resident CPC compartment and cardiomyocyte proliferation following clearance of senescent cells.

Although clearing senescent cells using a genetic approach is not feasible in humans, the senolytic (D+Q) pharmacological approach described here is clearly translatable and can be used to target a fundamental aging mechanism present in most tissues, including the heart. Pharmacologically eliminating senescent cells or inhibiting the production of their SASP has been shown to improve cardiovascular function [16, 24], physical function [20], enhance insulin sensitivity [11], prevent age-related bone loss [17], reduce frailty, and increase lifespan and health span [13, 20]. Indeed, D+Q administration over 3 months decreased senescent cell markers (TAF+ cells) in the media layer of the aorta from aged (24 months) and hypercholesterolemic mice, which was met with improved vasomotor function [16]. The approaches used in the present study, and thus far in previous studies, target global elimination of all senescent cells. Therefore, whether cell-specific elimination or local delivery of senolytics results in a greater effect in eliminating senescent cells and rejuvenating tissue regenerative capacity is yet to be determined.

Previous work has shown that senescent human primary preadipocytes as well as human umbilical vein endothelial cells (HUVECs) develop a SASP with aging, and conditioned medium from senescent human preadipocytes induced inflammation in healthy adipose tissue and preadipocytes [13]. Recently, the translational potential of using D+Q senolytics on human tissue was reported. Freshly isolated human omental adipose tissue obtained from obese individuals (45.7±8.3 years) contained increased numbers of senescent cells and increased secretion of SASP cytokines. Treatment with D+Q (1µM + 20µM) or vehicle for 48 hours decreased the number of TAF-, p16^Ink4a -^ and SA β-gal-positive cells, and decreased the secretion of key SASP components, PAI-1, GM-CSF, IL-6, IL-8 and MCP-1 [20]. Clearance of senescent human CPCs using D+Q in the present study also abrogated the SASP, and the deleterious impact of the SASP on impairing proliferation and inducing senescence to healthy, cycling competent CPCs in a co-culture environment was abolished with D+Q treatment. Thus D+Q can kill human senescent cells, including tissue specific stem/progenitor cells, and can attenuate the secretion of inflammatory cytokines associated with human age-related frailty [20].

The anthracycline doxorubicin (DOX) is an effective chemotherapeutic agent used to treat pediatric cancers, but is associated with progressive and dose-related cardiotoxicity that may not manifest itself until many years after treatment. In a juvenile mouse model of anthracycline-mediated late onset cardiotoxicity, Huang et al. [91] reported decreased number of CPCs, which showed impaired differentiation into the cardiomyocyte and endothelial lineage in the MI-border zone of animals that had been exposed to DOX as pups. DOX treatment also led to significantly less CPCs in the juvenile heart, and those that remained showed decreased proliferation and increased expression of p16^Ink4a^ at 12 days of age. Moreover, 72 hours of DOX treatment (10nM or 100nM) on isolated CPCs in vitro led to attenuated proliferation, reduced telomerase activity and induced expression of p16^Ink4a^ [91]. The present study showed that DOX treatment (0.2µM) for 24 hours to CPCs induced p16^Ink4a^, γH2AX and SA β-gal expression evidencing that CPCs had become senescent following DOX exposure. Future work should focus on the regenerative capacity of the heart following DOX exposure and whether senolytics could be used following DOX treatment and in later life to eliminate senescent cells and rejuvenate CPCs and the regenerative potential of the heart.

In conclusion, the present work demonstrates that in the aged and failing human heart a large majority of its resident CPCs are senescent and dysfunctional exhibiting impaired proliferation, clonogenicity and differentiation potential. Importantly, senescent CPCs have a SASP that can negatively affect cell behavior and render neighbouring cells into senescence. Elimination of senescent human CPCs *in vitro* attenuates the SASP and its deleterious effect. Senescent human CPCs are unable to contribute to repair and regeneration of the MI heart, in contrast to cycling competent, healthy human CPCs, which facilitate reparative processes in the damaged myocardium restoring LV function. The present work demonstrates approaches that eliminate senescent cells may be useful for treating age-related cardiac deterioration and rejuvenating the regenerative capacity of the aged heart. The next steps would be to determine whether senolytic approaches could be used in conjunction with cell therapy interventions to improve the environment (the ‘soil’) the cells (the ‘seeds’) are being transplanted into and the intrinsic reparative mechanistic processes that are compromised with age. Indeed, targeting senescent cells could also impact the potency of resident stem/progenitor populations in other aged organs. The present findings provide new insights into therapies that target senescent cells to prevent an age-related loss of regenerative capacity.

## Methods

### Human CPC isolation

Myocardial samples (~200mg each) were obtained from the right atrial appendage (n=119) of subjects with cardiovascular disease, aged 32-86 years. All subjects had given informed consented to take part in the study (NREC #08/ H1306/91). Samples were stored in saline on ice until ready to process (~1hr). All steps were performed at 4°C unless stated otherwise. Briefly, cardiac tissue was minced then digested with collagenase II (0.3mg/ml; Worthington Laboratories) in Dulbecco’s Modified Eagle’s Medium (DMEM; Sigma-Aldrich) at 37°C in a series of sequential digestions for 3 minutes each. Enzymatically released cells were filtered through a 40µm cell strainer (Becton Dickinson, BD) and collected in enzyme quenching media (DMEM + 10% FBS). The isolated cardiac cells were collected by centrifugation at 400g for 10 min, resuspended in incubation media (PBS, 0.5% BSA, 2 mM EDTA), and passed through an OptiPrep^™^ (Sigma-Aldrich) density gradient medium to remove large debris. This involved layering the cardiac cell population on top of 16% and 36% OptiPrep^™^:DMEM solutions and then centrifuging for 20 min at 800g with no brake. Larger debris collected at the bottom of the tube, allowing for selective retrieval of the small cell fraction from the upper layers. For the isolation of c-kit^pos^, CD45^neg^, CD31^neg^ CPCs, first the small cardiac cells were depleted of CD45^pos^ and CD31^pos^ cells by immunolabelling with anti-human CD45 and CD31 magnetic immunobeads (Miltenyi) diluted (1:10) in incubation media for 30 minutes at 4°C with agitation. After antibody binding, the CD45^pos^/CD31^pos^ cells were depleted from the preparation using magnetic activated cell sorting (MACS; Miltenyi). The elution cells, which were CD45^neg^, CD31^neg^ were then enriched for c-kit^pos^ cardiac cells through incubation with anti-human CD117 immunobeads (Miltenyi) (1:10) for 30 minutes at 4°C with agitation and again sorted using MACS according to the manufacturer’s instructions. Antibodies used for selection are detailed in **Supplementary Table 1**.

### Flow cytometry and FACS

The c-kit^pos^, CD45^neg^, CD31^neg^ human CPCs were analysed for hematopoietic, mesenchymal, and endothelial cell markers using a FACSCanto II™ flow cytometer (BD) as previously described [92]. The antibodies used for flow cytometry and FACS are reported in **Supplementary Table 1**. Respective isotype controls (Biolegend) were used as negative controls for all flow cytometry procedures. All antibodies were applied at 1:10 diluted in incubation media for 30 min at 4°C with agitation. Data were analysed using FlowJo^®^ software (FlowJo LLC).

SA–β-gal^pos^ CPC sorting was performed by fluorescent activated cell sorting (FACS) using an ImaGene Green C_12_FDG lacZ gene expression kit (Life Technologies, USA) according to the manufacturer’s instructions and protocols previously reported [93]. Briefly, freshly isolated CPCs were resuspended in 1ml of 33µM C_12_FDG β-gal diluted in pre-warmed DMEM/F12 media and incubated at 37°C for 30 minutes with agitation. The cell suspension was then sorted based upon SA–β-gal^pos^ fluorescence using FACS.

### Cell culture

Human CPCs were cultured on CELLstart™ (ThermoFisher Scientific)-coated cultureware in growth medium, which is composed of (1:1) Dulbecco’s MEM/Ham’s F12 (DMEM/F12; Sigma-Aldrich) and Neurobasal A medium (ThermoFisher Scientific) containing 10% Stemulate^®^ pooled human platelet lysate (Cook medical, USA), 2mM L-alanyl-L-glutamine (ThermoFisher Scientific), B27 and N2 supplements (ThermoFisher Scientific), leukemia inhibitory factor (LIF) (10ng/ml; Millipore), bFGF (10ng/ml; Peprotech), EGF (20ng/ml; Peprotech), insulin-transferrin-selenite (ITS; ThermoFisher Scientific), 1% penicillin/streptomycin (ThermoFisher Scientific), 0.1% gentamicin (ThermoFisher Scientific), and 0.1% Fungizone™ (ThermoFisher Scientific) in a humidified hypoxic incubator at 37°C, 5% CO_2_, 2% O_2_.

### Immunocytochemistry

Human CPCs were either freshly isolated or grown in culture and then directly cytospun onto poly-lysine-coated slides using a Shandon Cytospin 4 Cytocentrifuge (ThermoFisher Scientific). Slides were immediately fixed using Shandon Cell-Fixx (ThermoFisher Scientific). After fixation, cells were allowed to air dry before proceeding with immunostaining. To prepare cells for immunostaining, slides were incubated with 95% ethanol for 15 min at room temperature, washed with PBS, and in the case of intracellular staining, incubated in 0.1% Triton X-100:PBS at room temperature for 10 minutes. After washing with 0.1% tween:PBS and a 1 hr incubation in 10% donkey serum, cells were incubated overnight at 4°C with primary antibodies to p16^InkK4a^, γH2AX, or Ki67 (**Supplementary Table 2**) all applied at 1:50 in 0.1% Tween:PBS. Slides were then washed and incubated with corresponding Alexafluor^TM^ secondary antibodies (Life Technologies) for 1 hour at 37°C. The nuclear DNA of the cells was counterstained with 4’,6-diamidino-2-phenylindole (DAPI) (Sigma-Aldrich) at 1µg/ml and mounted using Vectasheild mounting media (Vector labs). The cells were viewed and imaged using an ApoTome fluorescent microscope (Zeiss) or A1 confocal microscope (Nikon). A minimum of 20 random fields of view at x20 magnification were used to determine the percentage of positively stained cells, expressed as a percent of total nuclei.

SA–β-gal (SABG) staining was performed using a SA–β-gal staining kit (Cell Signaling Technology) according to the manufacturer’s instructions and protocols previously reported [93]. In brief, human CPCs were either freshly isolated and directly cytospun onto poly-lysine-coated slides (ThermoFisher Scientific) or grown in culture and then fixed *in situ* using 2% paraformaldehyde: PBS (vol/vol) (Sigma-Aldrich) for 5 minutes at room temperature. Following fixation, cells were washed with PBS before being incubated in SABG activity solution (pH 6.0) at 37°C overnight. The enzymatic reaction was stopped by washing slides with ice-cold PBS and SABG staining was fixed with ice-cold methanol for 30s before mounting/visualising. Senescent CPCs were identified as blue-stained cells using light microscopy. A minimum of 10 images were taken at x4 magnification from random fields and the percentage of SA–β-gal cells were expressed as percentage of total nuclei.

### Q-FISH analysis

Freshly isolated human CPCs were cytospun and fixed as described above. To perform Q-FISH, slides were prepared using a telomere PNA FISH kit (Dako) according to the manufacturer’s instructions. In brief, slides were incubated in 95% ethanol for 5 min and washed in Tris-buffered saline (TBS; 2 min), fixed in formaldehyde (3.7%) in PBS (2 min), washed in TBS (2 x 5 min), and a pre-treatment solution applied at 1:2000 in TBS (10 min). Slides were washed again with TBS (2 x 5 min), dehydrated in cold ethanol series (70%, 80%, 95%, 2 min each), and air dried. The PNA hybridization mixture (10μl) containing Cy-3-conjugated (C_3_TA_2_)_3_ peptide nucleic acid (PNA) probe was added to the cells, a coverslip (18 × 18 mm) was then applied, followed by DNA denaturation (5 min at 80°C). After hybridization for 30 min, slides were incubated in wash solution with agitation (5 min at 65°C). Slides were 2% FBS counterstained with DAPI (Sigma-Aldrich) and mounted using Vectashield mounting media (Vector labs). All steps were performed at room temperature unless otherwise stated. Images were captured at 100x magnification on an A1 confocal microscope (Nikon). To prevent a possible selection bias, images were acquired ‘blindly’ *i.e*., cells were chosen solely on the basis of DAPI staining without checking the corresponding Cy-3 image. Telomere fluorescence signals were measured from 20 single cells per sample using ImageJ software (Fiji, USA). Cells and telomeres were identified through segmentation of the DAPI image and the Cy-3 image, respectively. The integrated fluorescence intensity for each telomere was calculated after correction for background, based on the values of the surrounding pixels. Telomere fluorescence values were converted into Kb by calibration with the L5178-Y and L5178Y-R mouse lymphocyte cell lines with known telomere lengths of 10.2 and 79.7 kb, respectively [94].

### Proliferation, clonogenicity, cardiosphere formation and cardiomyocyte differentiation assays *in vitro*

A BrdU incorporation assay (Roche) was used to determine cell proliferation, as previously described [95]. In brief, 2.5 x 10^3^ CPCs were plated in 24 well plates and serum-starved for 6 h in DMEM/F12 medium. Medium was then replaced with growth medium and BrdU was added, 1μg/ml every 8 hours. Cells were fixed after 24 hours and BrdU incorporation was assessed using a BrdU detection kit (Roche) according to the manufacturer’s instructions (**Supplementary Table 2**). Nuclei were counterstained with the DAPI (Sigma-Aldrich). Cells were viewed and imaged using an ApoTome fluorescent microscope (Zeiss). A minimum of 10 random fields of view at x20 magnification were analysed and number of BrdU-positive cells expressed as a percent of total nuclei.

Human CPCs (P2-P3) were serially diluted and deposited into CTS CELLstart pre-coated 96-well plates (Corning Inc., USA) at 1 cell per well, as described previously [83]. Individual cells were cultured in growth medium which was refreshed every 3 days. Wells containing CPC colonies were scored by bright-field microscopy after 7 days of culture to determine clonal efficiency. The clonogenicity of the human CPCs was expressed as the percent number of clones generated from the total number of single cells plated. A minimum of 3 plates were quantified per subject.

Cardiomyocyte differentiation was induced as previously described [81, 83]. Briefly, 5x10^4^ human CPCs were treated with oxytocin (100 nM; Sigma-Aldrich) for 72 h. Human CPCs were then transferred to a 10cm bacteriological dish containing LIF–deprived CPC growth medium to generate spheres. Spheres were transferred to laminin (1μg/ml; Sigma-Aldrich)-coated coverslips within 24 well plates and incubated with cardiomyogenic differentiation medium: alpha-MEM, 2% FCS, ascorbic acid (50μg/ml), dexamethasone (1 μM), beta-glycerol phosphate (10 mM), 1% penicillin-streptomycin, 0.1% gentamicin, and 0.1% Fungizone. Differentiating cultures were maintained for up to 14 d, replacing the medium containing specific growth factors on days 1 and 5. From days 1–4, cells were incubated with medium containing BMP-2 (10 ng/ml; Peprotech), BMP-4 (10 ng/ml; Peprotech), and TGF-β1 (5 ng/ml; Peprotech). From days 5 to 14, cells were incubated with medium containing 150 ng/ml Dkk-1 (150ng/ml; R&D). Medium was replaced every 3 days and cultures were fixed and analysed at 14 days. Cardiosphere formation was quantified from a minimum of 10 images per subject taken at x4 magnification from random fields. The number of cardiospheres were expressed per cm^2^. Cardiosphere size was determined from the same images by measuring the diameter of the spheres using ImageJ software (Fiji, USA). A minimum of 20 spheres per donor were measured. Cardiomyocyte differentiation was measured by immunostaining of differentiated cultures using Nkx2.5 (1:20, overnight 4°C) and α-sarcomeric actin (1:50, 1 hour 37°C) antibodies (**Supplementary Table 2**). Secondary antibodies were Alexa Fluor 488, or 594 (Life Technologies). Nuclei were counterstained with DAPI. A minimum of 5 images at x20 magnification from random fields were taken per subject for quantification of Nkx2.5^pos^ cells per total nuclei and fluorescent intensity of α-sarcomeric actin expression.

### Senescence induction, conditioned media, and multiplex SASP protein analysis

To induce senescence pharmacologically, cycling-competent CPCs were exposed to Rosiglitazone (0.1 µM; Calbiochem) or Doxorubicin (0.2 µM; Calbiochem) at ~70% confluence for 24 h. Medium was removed and cell cultures washed three times with PBS before replacing with fresh growth medium. Culture medium was refreshed every 3 days and CPCs were maintained for up to 35 days. Cells were analysed for SA-β-gal (as described above) and p16^INK4A^ expression over this time course by quantifying the total number of positive cells per total nuclei from a minimum of 5 random fields at 10x and 20x magnification respectively. Conditioned media (CM) were prepared by pre-washing cycling-competent or senescent CPC cultures (5x10^5^ cells) three times with PBS, then exposing them to RMPI 1640 containing 1 mM sodium pyruvate, 2 mM glutamine, MEM (minimum essential medium), vitamins, MEM non-essential amino acids, and antibiotics (ThermoFisher Scientific) for 24 h. CM was filtered through a 0.2 µM filter and stored at -20°C until ready for use or analysis. Luminex^®^ xMAP technology was used to quantify SASP factors in CM. Multiplexing analysis was performed using the Luminex 100 system (Luminex) by Eve Technologies using human multiplex kits (Millipore).

### PKH26 cell labelling

Human cycling-CPCs, Senescent-CPCs or c-kit^neg^ cells (5x10^5^) were harvested and washed in serum-free DMEM/F12 medium before being resuspended in 25µl diluent C (Sigma-Aldrich). An equal volume of PKH26 labelling solution (4µM; Sigma-Aldrich) was added to the cell suspension and mixed thoroughly. Cells were incubated for 2 minutes at room temperature in the dark, before adding an equal volume of DMEM/F12 + 10% FCS to halt the labelling reaction. Cells were then centrifuged and washed three times in PBS before transferring to a new Eppendorf tube and re-suspending in 15µl PBS for transplantation *in vivo*. Initial PKH26 labelling was validated using flow cytometry compared to unlabelled cells. PKH26 labelled cycling CPCs and c-kit^neg^ cells were also propagated *in vitro* to monitor label retention over culture passage (P1-P9). To quantify PKH26^pos^ labelling, cells were cytospun and counterstained with DAPI. The total number of positive cells per total nuclei were quantified from a minimum of 5 random fields at x20 magnification.

### Acute myocardial infarction model

The following experimental procedures were conducted in accordance with the regulations for animal testing, directed by the Home Office and stipulated under the Animals (Scientific Procedures) Act 1986. Non-obese diabetic (NOD), severe combined immunodeficiency (SCID) IL2Rgamma-c null mice [NSG; Charles River, UK] were used for all acute myocardial infarction studies. Mice were group housed in conditions of 22±1°C room temperature and 50±1% relative humidity on a 12-hour light-dark cycle. Standard laboratory mice chow and water were available *ad libitum*. Mice were 8-10 weeks old and were randomly assigned to each treatment group. Sample sizes were estimated according to power calculations performed on Mintab Software. Sham (n=5), Cycling-CPCs (n=7), Sen-CPCs (n=5), c-kit^neg^ cells (n=6)

Prior to echocardiography, mice were weighed and anaesthetised in an anaesthetic chamber at 0.5ml/min O_2_ and 4% Isoflurane. The depth of anaesthesia was assessed by a change in the breathing pattern of the mouse and the absence of the pinch withdrawal reflex. Once under deep anaesthesia, mice were placed onto an imaging station (Vevo, Netherlands). Anaesthesia was regulated to maintain heart rate between 400-500 beats per minute monitored by ECG recordings, and body temperature was maintained at 37 ± 0.5^°^C using a temperature probe. Mice were depilated and ultrasound gel was applied on to the vicinity of the chest (Aquasonic 100, Germany). Parasternal long and short axis images of the heart were taken using the Vevo 770 imaging system (Netherlands) and measurements taken using the B-mode and M-mode features. Ultrasounds were taken prior to left anterior descending coronary artery ligation (Baseline) and 1 and 4 weeks post-surgery. The following measurements were taken: interventricular septum thickness at end-systole/diastole (IVS_s/d_), left ventricular end diastolic volume/ left ventricular end systolic volume (LVEDV/LVESV), and left ventricular anterior wall thickness at end-systole/diastole (LVAW_s/d_). Ejection Fraction (EF), Stroke Volume (SV), and Fractional Shortening (FS) were measured manually using the left ventricular interventricular dimension (LVID) Trace tool and were also derived by long and short axis calculations.

Left anterior descending (LAD) coronary artery ligation was achieved as follows: once under deep anaesthesia, mice were intubated and attached to a mechanical ventilator (200ml stroke volume; 150 breaths per minute, Hugo Sachs Electronik, Germany), anaesthesia was reduced to 2% isoflurane at 0.5ml/min for the duration of the surgical procedure. Proceeding depilation, the thoracic area was sterilised and a midline incision along the skin overlying the sternum was performed to expose the muscle. A left-sided thoracotomy between the fourth and fifth rib was performed to expose the heart. Following the separation of the pericardial sac, the coronary artery was ligated using 6-0 silk suture (Ethicon, US) using the left atrium and aorta as entry and exit landmarks, respectively. The force of the suture was determined by the appearance of a color change in the left ventricle. Following the permanent ligation, the heart was assessed for atrial fibrillation for a 10 minute interval and was preceded by the closure of the thorax and the skin using 5-0 silk suture (Ethicon, US). In sham-treated mice, a permanent ligature was not tied around the LAD coronary artery. Immediately following MI, mice were injected intramyocardial with PKH26 labelled cycling-CPCs, Senescent-CPCs, or c-kit^neg^ cells (all 5x10^5^) delivered in a total volume of 15µl PBS, delivered across two sites at the border zone. Directly following MI+cell injection, mice were implanted with an osmotic pump (Alzet^®^, USA) loaded with a 0.2M solution of BrdU (MP Biomedicals), releasing the thymidine analogue for 14 days. Anaesthesia was then removed and mice were taken off mechanical ventilation after observing an independent breathing pattern dyssynchronous to that of the ventilator. Methadone analgesic was administered, Comfortan^®^ (1mg/kg; Dechra) i.m., and mice were kept in heat boxes overnight in a constant warm environment (27±1°C and a relative humidity of 50±5%). 24 and 48 hours following surgery, body weight and signs of pain were checked and post analgesic was administered as and when necessary. Following surgery, mice were housed singly caged with access to water and chow *ad libitum*. Echocardiography measurements were taken at baseline (BL), 7 days, and 28 days post-MI, after which mice were sacrificed.

### Senolytic drug treatment, viability, TUNEL staining *in vitro*

Cycling-competent human CPCs and Senescent-CPCs, induced by Doxorubicin (as described above), were plated in separate 24-well plates. Senescence was confirmed by SA-β-gal staining and the percentage of SA-β-gal^pos^ cells at day 0 was over 80% prior to senolytic drug treatment. After 3 days of either Dasatinib (LC Laboratories), Quercetin (Sigma-Aldrich,ST. Louse, MO), Navitoclax (Selleckchem, Houston, TX), Fisetin (Sigma-Aldrich, St. Louise, MO) at doses 0.5µM to 20µM or vehicle exposure, the number of Senescent-CPCs was quantified by SA-β-gal staining.

Cell viability was measured by crystal violet (CV) staining at day 0 then following 3 days of drug exposure. Briefly, cells were washed twice with PBS, fixed in methanol on ice for 10 minutes, and stained with 0.5% crystal violet (Sigma-Aldrich) in methanol for 15 minutes at room temperature. Cells were washed with deionized water and staining intensity was measured using a light microscope, with a minimum of 5 random fields of view at 4x magnification taken per well. The captured images were analyzed using ImageJ software (Fiji, USA).

Cell apoptosis was detected using the terminal deoxynucleotidyl transferase (TdT)-mediated dNTP nick end labelling (TUNEL) assay (Trevigen) according to the manufacturer’s instructions. Cells were cultured in 24-well plates as described previously and then treated with either vehicle or D+Q for 16 hours. To perform TUNEL staining, cells were incubated with proteinase K for 15 min at room temperature then a labelling buffer containing TdT enzyme, TdT dNTP, and Co^2+^ was applied for 1h at 37°C. Positive controls were incubated with TACS nuclease, while negative controls omitted the TdT enzyme from the labelling mix. After labelling, cells were washed in stop buffer for 5 min and were then incubated with Strep-Fluorescein in 0.05% Tween:PBS for 20 min at room temperature. After washing, cells were counterstained with DAPI (Sigma-Aldrich) and mounted using Vectasheild mounting media (Dako). The cells were viewed and imaged using an ApoTome fluorescent microscope (Zeiss). A minimum of 5 random fields of view at x20 magnification were analysed and number of TdT-positive cells expressed as a percent of total nuclei.

### Co-culture assays

CPCs were plated in the bottom well of 0.4µm pore Costar Transwell 24-well plates (Corning) and, once attached, were induced to senescence with Doxorubicin as described above and maintained for 30 days. Cycling-competent CPCs were plated at low density (200 cells/well) onto the Transwell membrane 1 day prior to co-culture. Cycling-competent CPCs were then combined with either senescent CPCs or cultured alone (control wells) and maintained for 7 days. At this time point, cultures were either fixed for analysis, conditioned media were collected, or senolytics were applied to transwells for 3 days to selectively clear senescent cells (as described above). After senolytic clearance, a sub-set of wells were fixed and stained for SA-β-gal (as described previously) to confirm effective clearance. Cycling-competent CPCs were maintained for a further 7 days, at which point conditioned media were collected for luminex analysis (as described above) and cultures fixed for analysis.

### INK-ATTAC mice and drug treatments

The following experimental procedures were conducted in accordance with Mayo Clinic Institutional Animal Care and Use Committee (IACUC) guidelines. Both stocks of *INK-ATTAC* and C57BL/6 wild type mice were bred and aged at Mayo Clinic. The generation and characterization of *INK-ATTAC* mice has been described previously [9, 11]. Mice were housed in ventilated cages and maintained in a pathogen-free, accredited facility under a 12 h light–dark cycle with constant temperature (23°C) and access to food (standard mouse diet, Lab Diet 5053, St. Louis, MO) and water *ad libitum*. 24-32 month old mice were randomly assigned to treatment groups and injected intraperitoneally (i.p.) with either vehicle or AP20187 (B/B homodimerizer, Clontech, 10 mg AP20187/kg body mass) or administered by oral gavage vehicle (66% Phosal-50PG, 10% ethanol, and 30% PEG-400) or senolytic drugs (5mg/kg Dasatinib + 50mg/kg Quercetin diluted in vehicle) for 3 consecutive days every 2 weeks for 2 months as shown in Fig 6a. This dosing regime was chosen because it has been shown to be effective at clearing senescent cells in chronologically aged *INK-ATTAC* mice in previous studies [11, 16]. Another group of *INK-ATTAC* mice aged 3 months (n=10) and 22 months (n=10) were randomly assigned to treatment groups and injected intraperitoneally (i.p.) with either vehicle or AP20187 for 2 consecutive days every 2 weeks for 2 months as shown in Supplementary Fig 10c. These mice were injected (i.p) with EdU (123 mgkg^-1^) 2 hours prior to sacrifice.

### Tissue collection, immunohistochemistry, and confocal imaging

Mice were sacrificed at either 4 days or 28 days after MI with the hearts arrested in diastole with cadmium chloride solution (Sigma), removed, and embedded in OCT compound (Tissue-Tek) at ^_^80°C before preparing 10μm sections. To identify transplanted cells, transverse sections were identified by PKH26 fluorescent labelling (Sigma-Aldrich) and nuclei counterstained with DAPI. Sections were assessed for co-localisation of PKH26 labelling and myocytes or vascular structures using antibodies against α-sarcomeric actin (cardiomyocytes; 1:250 dilution, 1hr at 37°C) or vWF (endothelial cells; 1:250 dilution, 1 hr at 37°C) respectively. Images were taken using a confocal microscope (Nikon A1) at x63 magnification. To quantify PKH26-cell engraftment, the total number of PKH26^pos^ cells per total nuclei were quantified from a minimum of 5 random fields at x20 magnification. To measure scar size, sections were stained with Haematoxylin Van Gieson (HVG, Fisher Scientific, UK) using standard procedures. Fibrosis was visualised using a light microscope (Zeiss, Axioscope MTB2004) connected to Axiovision software (Zeiss) where sections were captured at x5 magnification and stitched to reconstruct an image of a section of the whole heart. Analysis was performed using Image J software (NIH). Data were acquired by measuring the area of fibrosis (red) and the area of left ventricle and were expressed as a percentage mean ± SEM. (Zeiss Apotome fluorescent microscope, Oberkochen, Germany). Proliferating cells were identified using antibodies against BrdU (Roche) (Supplementary Table 1) and nuclei counterstained with DAPI. A minimum of 5 random fields of view at x20 magnification were analysed and data expressed as percent of BrdU-positive cells per total nuclei. Sections were also assessed for number of newly formed myocytes using antibodies against BrdU (Roche) and α-sarcomeric actin (1:250 dilution, 1hr at 37°C). A minimum of 5 random fields of view were taken at the border zone using at x63 magnification per animal. The number of BrdU-positive cardiomyocytes were expressed as a percent of total cardiomyocyte nuclei. Newly formed vascular structures were detected by staining for BrdU (Roche) and vWF (1:250 dilution, 1hr at 37°C). A minimum of 5 random fields of view at x20 magnification were analysed at the border zone and number of BrdU-positive capillaries (1-2 nuclei) expressed as a percent of total capillaries. In all cases sections were imaged and analysed using an A1 confocal microscope (Nikon). All antibodies used in immunostaining are listed in **Supplementary Table 2**.

INK-ATTAC and wild-type C57BL/6 mice were sacrificed 4 days after the last dose of the last course of treatment with AP20187 or senolytic drugs. Hearts were explanted and cut into two parts. One half was fixed in 4% formaldehyde for 24 hrs and embedded in paraffin. The other half was snap-frozen in liquid nitrogen for subsequent RT–qPCR analyses as described below. 10μm transverse heart sections were cut on a microtome (Leica) and mounted onto microscope slides. Antigen retrieval was achieved using Target Retrieval Solution, Citrate pH 6 (DAKO). Sections were stained for c-kit (1:500 dilution, overnight at 4°C), Sca-1 (1:500 dilution, overnight at 4°C), CD45 (1:1000 dilution, 1 hr at 37°C), CD34 (1:1000 dilution, 1 hr at 37°C), CD31 (1:1000 dilution, 1 hr at 37°C), Ki67 (1:500 dilution, 1 hr at 37°C) and αsarcomeric actin (1:100 dilution, 1 hr at 37°C). Secondary antibodies were Alexa Fluor 488, 594 or 633 (Life Technologies). The nuclei were counterstained with DAPI) (Sigma-Aldrich). Sections were mounted in Vectashield (Vector labs) and analysed and scanned using confocal microscopy (A1 confocal, Nikon). CPCs were identified as being Sca-1- and c-kit-positive, and CD45-, CD34-, CD31-negative. Proliferating CPCs were identified as Sca-1- and c-kit-positive and Ki67-positive. Proliferating cardiomyocytes were identified as Ki67-positive or EdU-positive and α-sarcomeric actin positive. CPCs were counted in a minimum of 10 random fields of view at x40 magnification and the number of CPCs expressed as a percent of total nuclei. Proliferating cardiomyocytes were counted across random fields of view of 3 cross sections at x63 magnification. A total of 1500 cardiomyocytes were counted per animal per section and the number that were Ki67- or EdU-positive expressed as percentage of cardiomyocytes. Cardiomyocyte diameters were measured across the nucleus on transverse sections of α-sarcomeric actin-positive stained cardiomyocytes using ImageJ software. A minimum of 60 cardiomyocytes were measured per animal. LV fibrosis was detected using HVG staining as described above. All antibodies used in immunostaining are listed in **Supplementary Table 2**.

### Real-time quantitative polymerase chain reaction (RT–qPCR)

To measure human gene expression, total cellular RNA was extracted and reverse transcription was performed as described previously [92]. RT-qPCR reactions were run with SYBR Green PCR Master Mix (Bio-Rad) and primers (IDT) (**Supplementary Table 3**). Human *GAPDH*, *B-actin*, and *TBP* were used as reference genes. qPCR conditions were as follows: 95°C for 5 min, 40 × (95°C for 15s, 60°C for 30s, and 72°C for 30s). All reactions were run in triplicate on a CFX-connect Real-Time PCR System (Bio-Rad) and analyzed using Graphpad software.

From mouse hearts, total RNA was extracted using Trizol (Thermo Fisher Scientific) and reverse transcription was performed using an M-MLV reverse transcriptase kit as described previously [13]. To measure total *p16^Ink4a^* expression, RT-qPCR reactions were performed with TaqMan (Applied Biosystems, Carlsbad, CA) Fast Advanced Master Mix and TaqMan primer- probe gene assay (Thermo Fisher Scientific). Mouse TATA-binding protein (*TBP)* was used as a reference gene and all reactions were run in triplicate on an ABI Prism 7500 fast Real Time System (Applied Biosystems, Carlsbad, CA) and analyzed using Graphpad software.

### Statistical analysis

Data are reported as mean ± SEM or mean ± SD. The data displayed normal variance. The experiments were not randomized, except for the *in vivo* animal studies as described above. The investigators were blinded to allocation during experiments and outcome assessment. Significance between 2 groups was determined by Student’s *t-*test and in multiple comparisons by the analysis of variance (ANOVA) using GraphPad Prism (GraphPad Software). In the event that ANOVA justified *post-hoc* comparisons between group means, these were conducted using Tukey’s multiple-comparisons test. *P*<0.05 was considered significant.

## ACKNOWLEDGEMENTS

We acknowledge Thomas Theologou and Mark Field from Liverpool Heart & Chest Hospital, Institute for Cardiovascular Medicine and Science (ICMS), Liverpool, UK for giving us some myocardial samples. We thank Carl Hobbs (King’s College London) for tissue processing, histology, and microscopic analysis assistance. Confocal microscopy was carried out in the Nikon Imaging Centre, King’s College London.

## AUTHOR CONTRIBUTIONS

F.C.L performed the isolation and characterisation of CPCs from subjects’ myocardial samples, functional CPC assays, expression and secretion of SASP factors, senolytic clearance of senescent cells *in vitro*, immunohistochemical and confocal microscopy analysis of mouse cardiac sections. P.J.R performed the myocardial infarction-regeneration mouse surgery model and echocardiography. E.D-V performed the characterisation of senescent-CPCs for SASP factors and the effect of SASP factors on CPC fate. T.S.T performed isolation and characterisation of CPCs for p16^Ink4a^ expression. B.J.C analysed mouse cardiac sections using immunohistochemical and confocal microscopy. J.E.C oversaw and assisted with the myocardial infarction-regeneration mouse surgery model and echocardiography. P.P.P and W.A provided myocardial samples and contributed to experimental design. D.T contributed to experimental design. T.T and J.L.K contributed the senolytic drugs, performed the genetic and pharmacological elimination of senescent cells in mice, contributed to experimental design, and directed the elimination of senescent cells aspect of the study. L.P carried out the EdU INK-ATTAC experiments. G.M.E-H directed and supervised all aspects of the study, and wrote the manuscript with input from F.C.L, P.J.R, D.T, T.T., and J.L.K. All authors reviewed the manuscript.

## List of Supplementary Materials

Tables S1 – S3

Figures S1 – S10

